# The Potential of CCA-associated Bacteria to Fight Antimicrobial-Resistant Pathogens: a Genomic Survey

**DOI:** 10.1101/2024.11.26.625565

**Authors:** Diego Lera-Lozano, Jordan S. Ruiz-Toquica, Matthew W. Holt, Elizabeth K. Jones, Kim B. Ritchie, Mónica Medina, Raúl A. González-Pech

## Abstract

The global rise of antimicrobial resistance has intensified efforts in bioprospecting, with researchers increasingly exploring unique marine environments for novel antimicrobials. In line with this trend, our study focused on bacteria isolated from the unique microbiome of crustose coralline algae (CCA), which has yet to be investigated for antimicrobial discovery. In the present work, bacteria were isolated from the CCA *Hydrolithon boergesenii*. After running antimicrobial assays against antibiotic-resistant human and marine pathogens, three isolates were selected for genome sequencing using the Oxford Nanopore technology. Genome mining of the high-quality assemblies revealed 100 putative Biosynthetic Gene Clusters (BGCs) across the three genomes. Further analysis uncovered BGCs potentially involved in the biosynthesis of novel antimicrobial compounds. Our study provides valuable resources for future research aimed at the discovery of novel antimicrobials, particularly in the face of the antibiotic-resistance global crisis and highlights the potential of specialized marine environments like CCA biofilms.

## INTRODUCTION

Antimicrobial resistance poses a critical threat to public health globally, with estimates suggesting that by 2050, it could claim up to 10 million lives annually and cause substantial economic losses, impacting healthcare and human development [1][2]. In response to this growing crisis, the urgent need for new antimicrobials, particularly novel structural classes that can circumvent existing resistance mechanisms, has become paramount [3]. Considering this, bioprospecting efforts have intensified in pursuit of new antimicrobial compounds with the potential to target antimicrobial-resistant pathogens.

Over the past century, scientists have identified natural products with a wide variety of use cases, including agricultural implementations (e.g., pesticides), nutraceuticals (e.g., dietary supplements), and notably, pharmaceuticals (e.g., antibiotics). Bioprospecting has been critical in these discoveries. In particular, bioactive secondary metabolites have been identified from bacteria living in environments with a high microbial diversity due to their broad metabolic spectra [4]. For example, the commonly used antibiotic vancomycin was originally recovered from a soil bacterium [5]. Similarly, the antiviral scytovirin, used to treat HIV, was derived from the freshwater cyanobacterium *Scytonema varium* [6]. Bacteria living in marine sediments contributed to the establishment of the antitumor antibiotics marinomycins [7].

Coral reefs are composed of several environmental niches that support a diverse array of both macro- and microorganisms. Recent research has revealed that these ecosystems harbor a rich microbial community compared to many other environments [8]. This microbial richness is characterized by substantial heterogeneity in community composition and metabolic potential across different microenvironments within a single coral reef [9]. Microbial communities associated with reefs have been identified as promising sources of antimicrobial compounds. For example, an initial screening of culturable bacterial strains from the elkhorn horal (*Acropora palmata*) revealed that 20% exhibited antibiotic activity [10]. Additionally, coral-associated bacteria, like *Pseudovibrio* sp. P12, have yielded compounds such as tropodithietic acid, which effectively inhibits various coral pathogens resistant to commercial antibiotics [11].

While reef organisms like sponges and soft corals have provided notable antibiotics including halichondrin B and eleutherobin, respectively [12][13], microorganisms associated with other reef organisms have been relatively underexplored for novel antimicrobials. A prime example are crustose coralline algae (CCA). CCAs host diverse bacterial communities with significant ecological and biotechnological potential [14][15][16]. These bacteria produce unique metabolites that contribute to host defense and health. While some CCA-associated bacteria like *Propionibacterium* and *Planomicrobium* show antimicrobial activity [17][18][19], research has primarily focused on their role in coral larval settlement [20][21] and antifouling properties [22]. The rich microbial diversity and metabolite production in CCA microenvironments offer promising opportunities for novel antimicrobial discovery.

Biosynthetic gene clusters (BGCs) play a crucial role in microbial adaptation and survival [23]. They regulate the production of secondary metabolites, i.e., compounds not directly involved in the growth or reproduction of microorganisms but essential for their environmental interactions, such as preventing overgrowth by other microbes [24]. Their presence might be especially important in environments with intricate microbial interactions, such as CCA biofilms. Because of the diverse roles of secondary metabolites, BGCs can be implicated in the synthesis of a wide array of bioactive compounds, with antimicrobials among the most prominent [24]. Consequently, the identification of BGCs in bacterial genomes is an integral part of the bioprospecting process, particularly when the product responsible for antimicrobial traits remains unknown [24].

In this study, CCA-associated bacteria were isolated and tested for antimicrobial properties against antibiotic-resistant and other human and marine pathogens. Selected isolates were subjected to whole-genome sequencing to explore features that might explain their antimicrobial phenotypes. Several putative BGCs were identified, some potentially involved in the production of novel antimicrobial compounds. This study sets the foundation for future research aimed at discovering new antimicrobials and highlights the need to target niche marine environments, such as CCA, for bioprospecting.

## MATERIALS & METHODS

### Sample Collection, Bacteria Isolation, and Culture Conditions

The CCA *H. boergesenii* was isolated from coral rubble at 8 m at the Veradero Reef in the Cartagena Bay, Colombia, on March 7th, 2023. CCA fragments were stored in sterile 15 ml screw-capped tubes and swabbed within 3 h of collection using sterile cotton swabs, followed by culture onto marine agar (Difco) plates. Bacteria were subcultured to purification and stored in 96-well plates in 20% glycerol diluted in marine broth. CCA were taxonomically identified via microscopy.

### Screening for Antimicrobial Activity

Each of the bacterial isolates were analyzed for antibacterial properties against several human and marine pathogens using an agar-overlay assay method. Culture libraries from 96-well plates were replica plated onto sterile marine agar and allowed to grow for 6 days. Cultures were then exposed to UV radiation for one hour followed by overlay with 0.8% soft agar containing one of the following pathogenetic test strains: *Escherichia coli* (EC), *Bacillus subtilis* (BS), Vancomycin resistant *Enterococcus* (VRE), *Enterococcus* (ENT), Methicillin-resistant *Staphylococcus aureus* (MRSA), Methicillin-sensitive *Staphylococcus aureus* (MSSA), *Vibrio vulnificus*, *Vibrio shiloi*, *Vibrio parahaemolyticus*, and *Serratia marcescens* PDL100. Zones of growth inhibition were measured in mm after 2 days. Test strains (shown in Table 1) were grown overnight at 37 °C (human pathogens) or 25 °C (marine pathogens) in 5 mL of strain-specific culture broth. Aliquots of each broth culture were inoculated into 0.8% agar containing marine broth, Luria broth or tryptic soy broth, depending on the test strain. CCA library plates were overlaid with approximately 10 mL of inoculated agar. Following overnight growth, plates were analyzed for antibacterial activity and zones of inhibition were measured in millimeters.

**TABLE 1.**
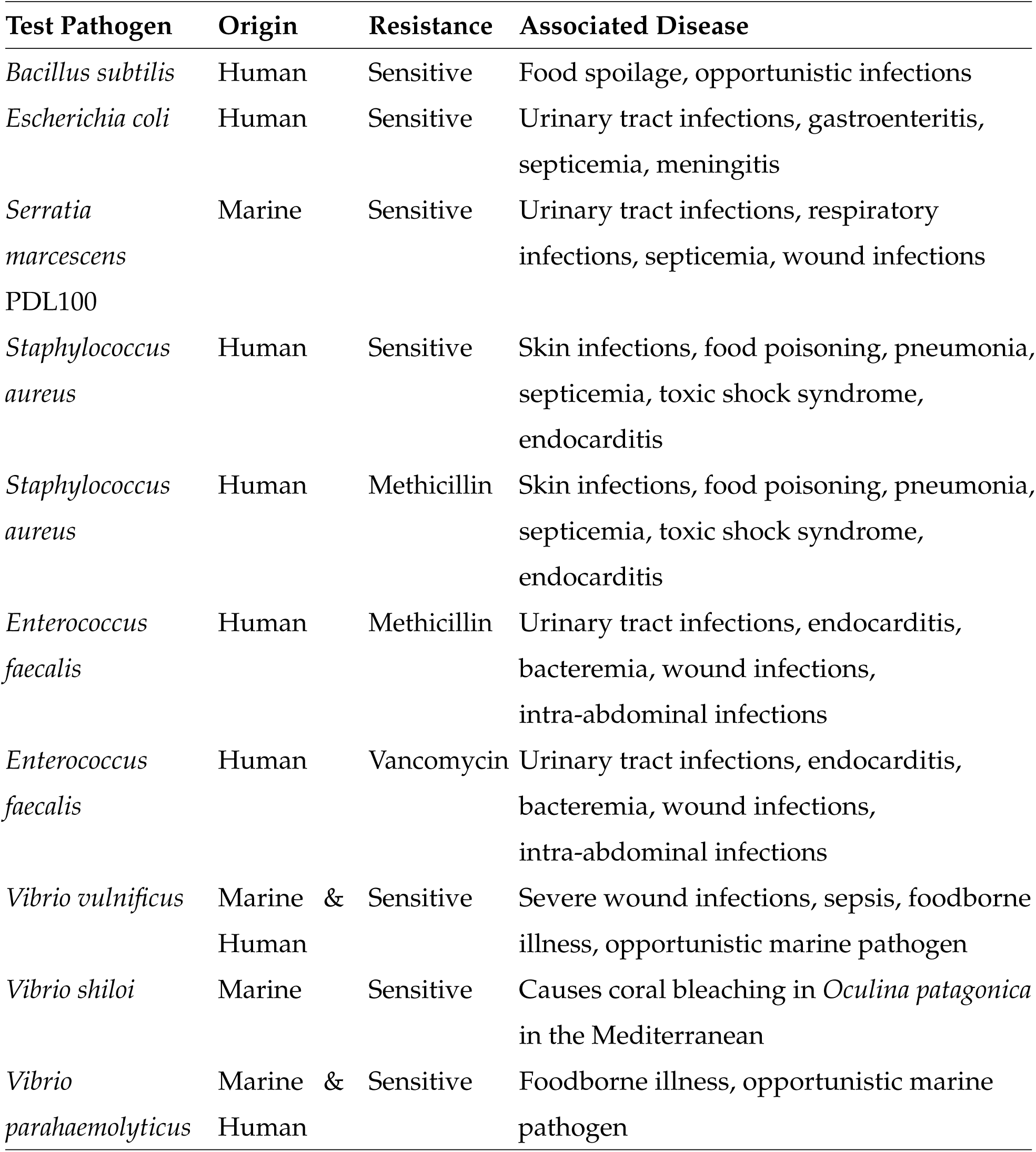
Human and marine pathogen test strains used in antagonistic assays, their antibiotic resistance profiles, and associated diseases.

| Test Pathogen | Origin | Resistance | Associated Disease |
| --- | --- | --- | --- |
| *Bacillus subtilis* | Human | Sensitive | Food spoilage, opportunistic infections |
| *Escherichia coli* | Human | Sensitive | Urinary tract infections, gastroenteritis, septicemia, meningitis |
| *Serratia marcescens*<br>PDL100 | Marine | Sensitive | Urinary tract infections, respiratory infections, septicemia, wound infections |
| *Staphylococcus aureus* | Human | Sensitive | Skin infections, food poisoning, pneumonia, septicemia, toxic shock syndrome, endocarditis |
| *Staphylococcus aureus* | Human | Methicillin | Skin infections, food poisoning, pneumonia, septicemia, toxic shock syndrome, endocarditis |
| *Enterococcus faecalis* | Human | Methicillin | Urinary tract infections, endocarditis, bacteremia, wound infections, intra-abdominal infections |
| *Enterococcus faecalis* | Human | Vancomycin | Urinary tract infections, endocarditis, bacteremia, wound infections, intra-abdominal infections |
| *Vibrio vulnificus* | Marine & Human | Sensitive | Severe wound infections, sepsis, foodborne illness, opportunistic marine pathogen |
| *Vibrio shiloi* | Marine | Sensitive | Causes coral bleaching in *Oculina patagonica* in the Mediterranean |
| *Vibrio parahaemolyticus* | Marine & Human | Sensitive | Foodborne illness, opportunistic marine pathogen |

### Genomic DNA Extraction and Sequencing

Genomic DNA (gDNA) was extracted from 24 h-marine agar cultures using the DNeasy Powersoil® Pro Kit (Qiagen, USA) with modifications to the cell lysis steps. Briefly, two to three colonies of each isolate were picked and suspended in 500 µl into the spun PowerBead Pro tubes and mixed with 800 µL of the CD1 solution. Tubes were then placed in a bead-beater vortex and shaken as follows: 1) 5 min at 13,000 rpm, 2) a 2 min pause, 3) incubation for 1 min at 4°C, 4) 5 min at 13,000 rpm again, and 5) a final incubation at 4°C for 2 min. After this step, we followed the manufacturer’s instructions. DNA was resuspended in 80 µL of elution buffer and quantified and quality-evaluated by electrophoresis in 0.8% agarose gels with 0.5 X TBE buffer (45 mM Tris-borate, 45 mM boric acid, 1 mM EDTA, pH 8.2) and using a Qubit fluorometer (Invitrogen™). The extracted DNA was further cleaned using a DNA Clean & Concentrator-5 kit (Zymo Research, Irvine, CA, USA) for high-purity absorbance ratios (260/280 = 1.7 and 260/230 = 2.2). DNA was then stored at -20°C until whole-genome sequencing on an Oxford Nanopore MinION Mk1B. Briefly, high-molecular-weight gDNA libraries were prepared using the Native Barcoding Kit 24 V14 (SQK-NBD114.24, Oxford, UK). Up to 400 ng of gDNA from each isolate were used to achieve equimolar concentration. Then, gDNA end-repair and ligation of native barcodes and adapters were done following manufacture’s protocol instructions. The libraries were then pooled and loaded into the primed R10.4.1 flow cell. MinKNOW v23.07.15 was used to monitor sequencing and for demultiplexing. Base calling was conducted separately using Dorado v0.4.3 [25] implementing the high-accuracy model dna_r10.4.1_e8.2_400bps_sup@v4.2.0 (optimized for Duplex pairs) with default settings other than disabling CUDA acceleration.

### Read Processing and Genome Assembly

Quality of raw reads was assessed using FastQC v0.12.0 [26]. Adapters were removed and reads were trimmed using chopper v0.7.0 [27], removing 100 bp from both 5’ and 3’ ends. Genome size and sequencing depth for each isolate were estimated using Jellyfish v2.3.0 [28] with k = 25 and GenomeScope 2.0 [29]. De novo genome assembly was done using Flye v2.9.2 [30], followed by circularization of complete molecules using Circlator v1.5.5 [31]. Quality of the final assemblies was assessed using CheckM v1.2.2 [32]. To determine whether contigs were of chromosomal or plasmid origin, we aligned each contig against the NCBI nucleotide database using BLASTn v2.16.0 [33]. Taxonomic assignment was done using GTDB-tk v2.3.2 [34] and the GTDB database release 207 [35].

### Gene Prediction and Functional Annotation

Genome annotation was done using prokka v1.14.5 [36] at default settings. Metabolic pathway analysis was done by first annotating predicted proteins with KEGG orthology using the blastKOALA web service [37] and then reconstructing metabolic pathways using the online KEGG mapper [38]. COGclassifier v1.0.5 [39] was used to assign predicted proteins into clusters of genes (COG) with annotated gene functions. COGs were also predicted from the genome of *Escherichia coli* strain K-12 substrain MG1655 (NCBI RefSeq Accession GCF_904425475) as a reference of a bacterium with little to none known secondary metabolite production [40]. Genes implicated in the production of tetrabromopyrrole (TBP)[41] were manually searched in the prokka annotations.

### Biosynthetic Gene Clusters Identification and Network Analysis

Biosynthetic gene clusters (BGCs) were identified using antiSMASH v7.0.0 [42], DeepBGC v0.1.29 [43], and GECCO v0.9.10 [44]. Circular visualization of genome assemblies and features was done in R using shinyCircos v2.0 [45]. BiG-SCAPE v1.1.9 [46] and the MIBiG database release 2.1 inclusion [47] were used for gene similarity network analyses based on the BGCs identified by antiSMASH, DeepBGC, and GECCO. One run was done using all BGCs as input and a gene similarity cutoff of 0.7; networks resulting from this run were considered *conservative*. *Less-conservative* networks were obtained from a second run with the same input but a similarity cutoff of 0.3. Clusters resulting from the similarity network analyses were visualized using Cytoscape v3.7.0 [48].

## RESULTS

### CCA-Associated Bacteria Inhibit Growth of Human Pathogens

The 14 bacterial strains isolated from the CCA *Hydrolithon boergesenii* were found to produce several zones of inhibition against the test pathogens with the results listed in Table 2. Three isolates in particular displayed the capacity to inhibit three or more of the test strains and were selected for subsequent genomic characterization.

**TABLE 2.**
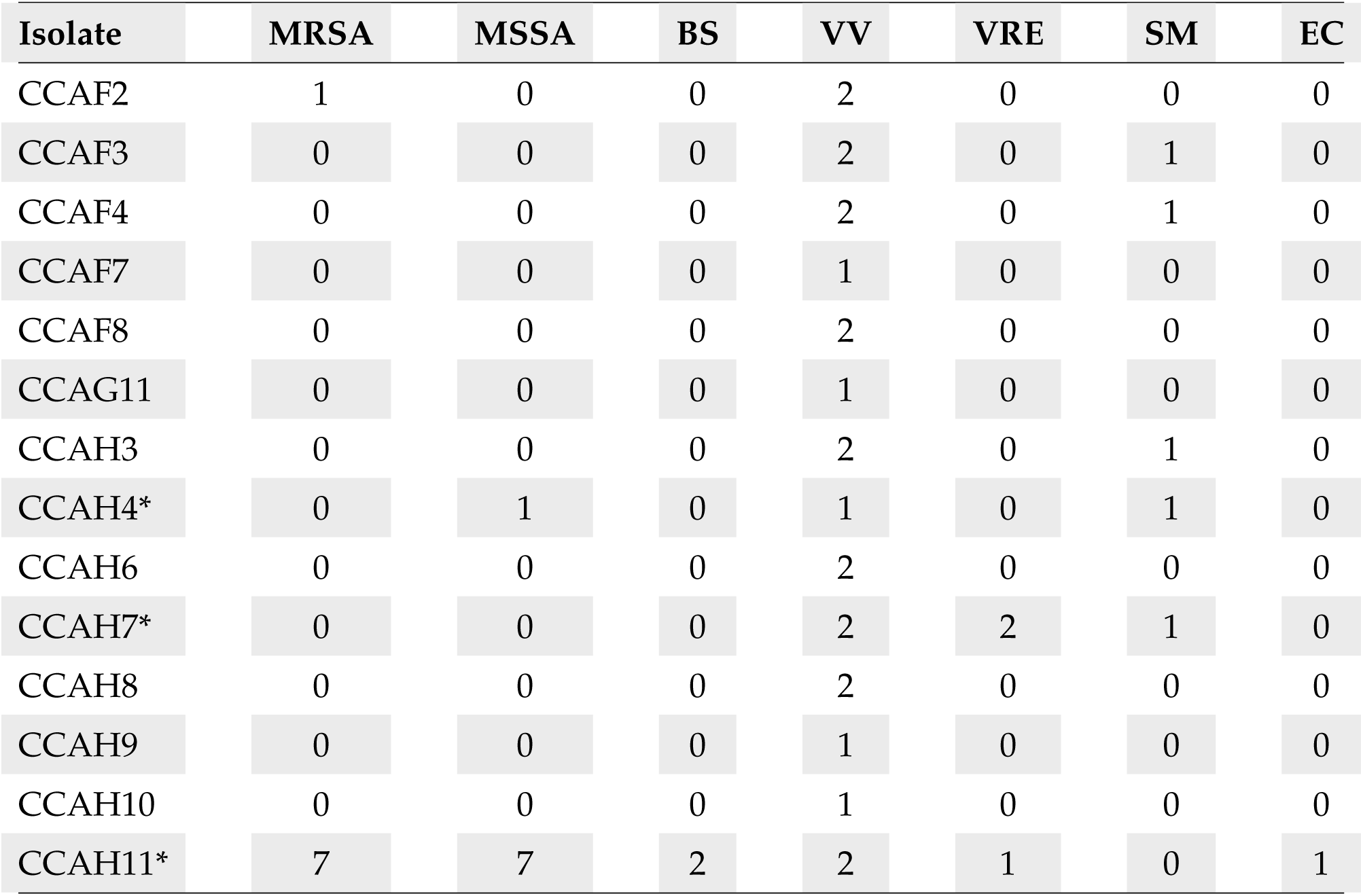
Results of antagonistic assays displaying the inhibitory activity of isolates against various pathogens (in millimeters). An asterisk (*) indicates that the corresponding isolate was selected for sequencing. MRSA: Methicillin Resistant *Staphylococcus aureus*; MSSA: Methicillin Sensitive *Staphylococcus aureus*; BS: *Bacillus subtilis*; VV: *Vibrio vulnificus*; SM: *Serratia marcescens* PDL100; VRE: Vancomycin Resistant *Enterococcus faecalis*; EC: *Escherichia coli*.

### Genomic Characterization of CCA Bacterial Isolates

Taxonomic assignment resolved the bacterial isolates to the species level as listed in Table 3. Based on an Average Nucleotide Identity (ANI) cutoff of *≥*95%, strain CCAH7 was identified as *Pseudovibrio denitrificans* (ANI = 95.29%), CCAH11 as *Pseudoalteromonas elyakovii* (ANI = 97.29%), and CCAH4 as *Rossellomorea marisflavi* (ANI = 98.92%). Genomic features of the three bacterial isolates are summarized in Table 3 and Fig. 1. The genomes were assembled into 3-4 contigs with N50 lengths ranging from 3.16 to 4.9 Mbp and high completeness (98.64%-100%). All assembled contigs in *P. denitrificans* CCAH7 consisted of entire circular molecules, as did one plasmid contig of *P. elyakovii CCAH11*; no circular molecules were recovered for *R. marisflavi* CCAH4 as listed in Table 3. The predicted protein-coding sequences (CDS) ranged between 4,435 and 5,331 per isolate.

**FIG 1.**
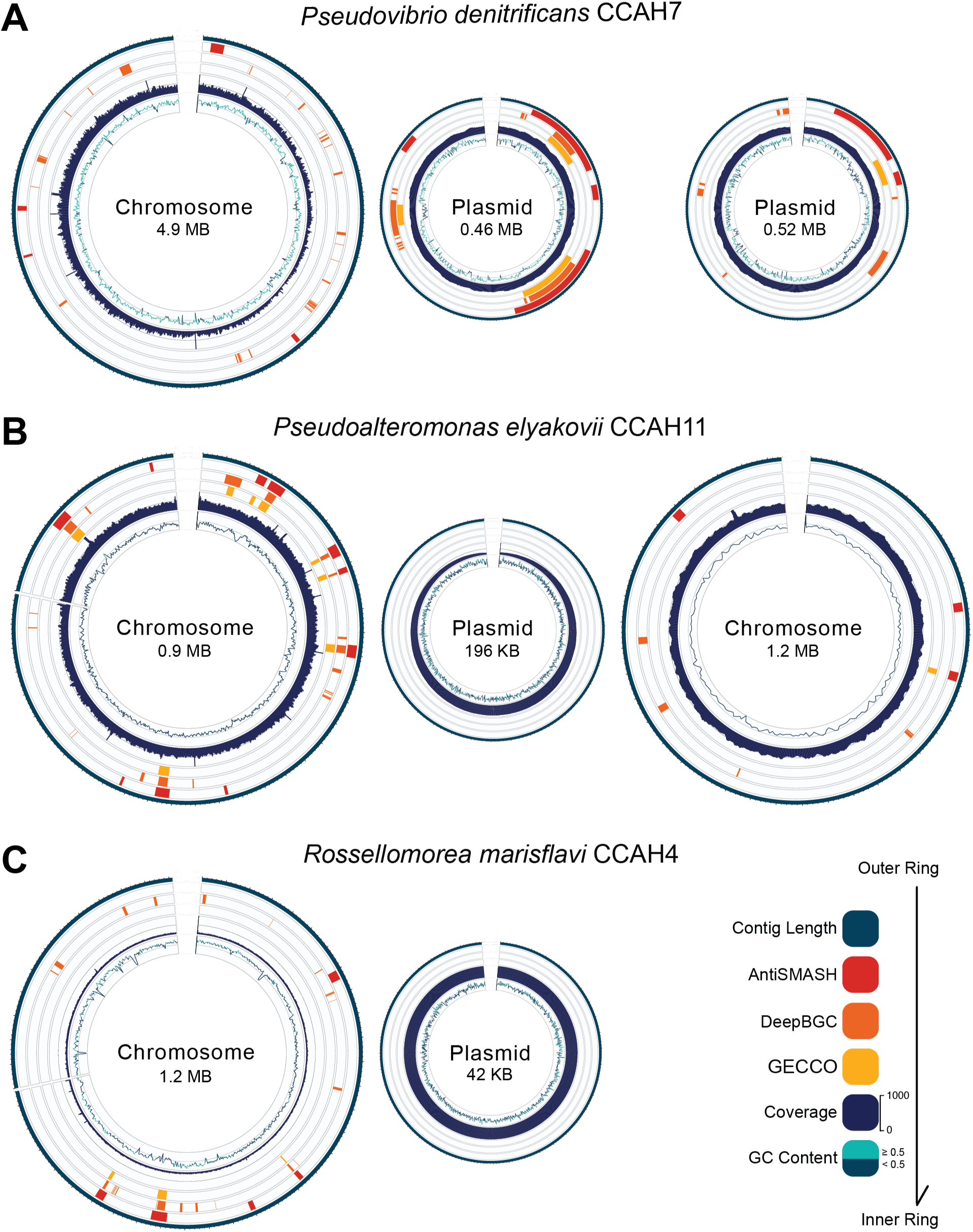
Circular genome representations of *Pseudovibrio denitrificans* **(A)**, *Pseudoalteromonas elyakovii* **(B)**, and *Rossellomorea marisflavi* **(C)**, depicting the distribution of biosynthetic gene clusters (BGCs) across chromosomes and plasmids. Each representation displays, from outer to inner rings, contig length, BGCs identified by AntiSMASH, DeepBGC, and GECCO, sequencing coverage, and GC content, following the bottom right color code.

**TABLE 3.**
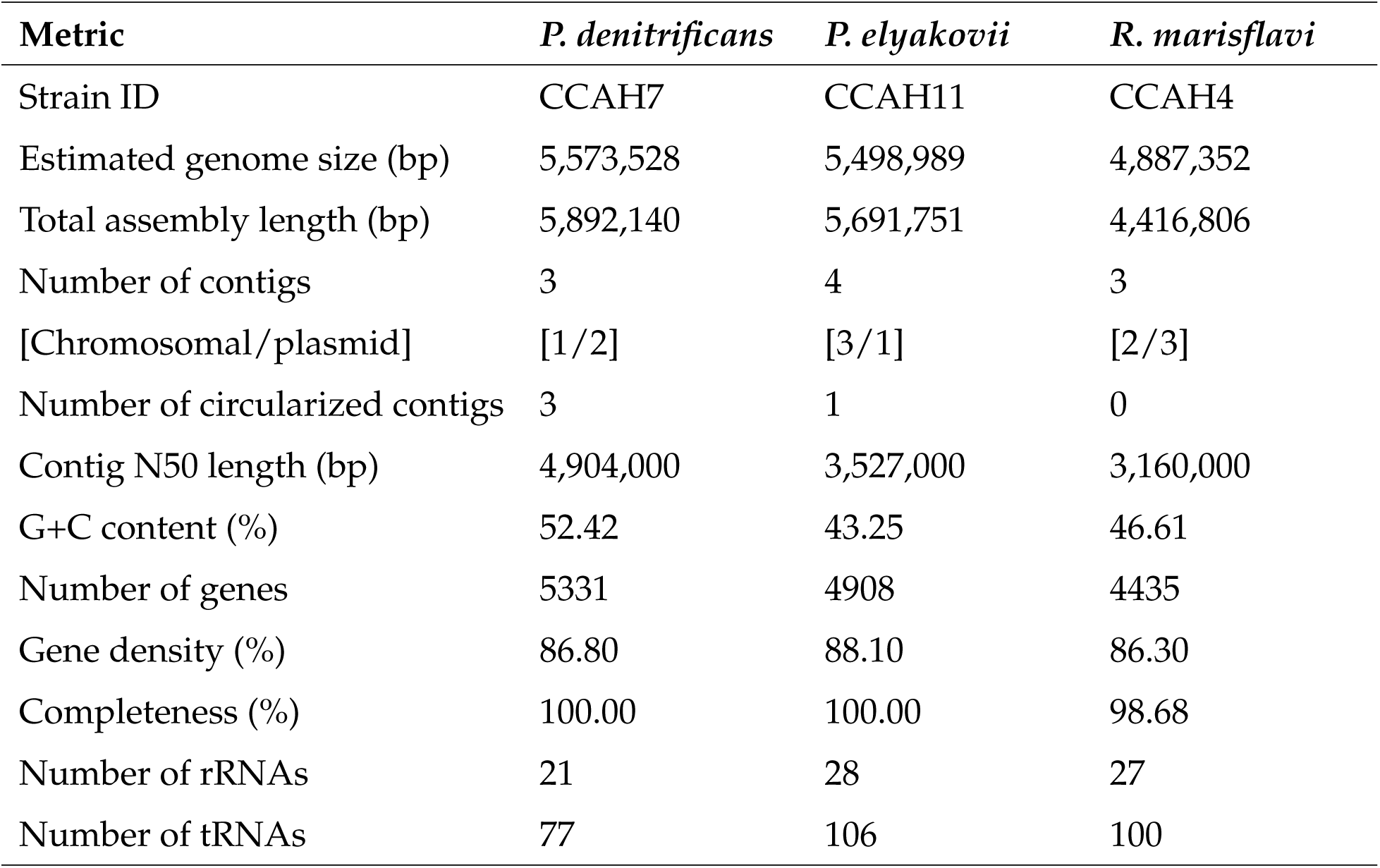
Summary of genome statistics for the sequenced bacterial isolates.

| **Metric** | ***P. denitrificans*** | ***P. elyakovii*** | ***R. marisflavi*** |
| --- | --- | --- | --- |
| Strain ID | CCAH7 | CCAH11 | CCAH4 |
| Estimated genome size (bp) | 5,573,528 | 5,498,989 | 4,887,352 |
| Total assembly length (bp) | 5,892,140 | 5,691,751 | 4,416,806 |
| Number of contigs | 3 | 4 | 3 |
| [Chromosomal/plasmid] | [1/2] | [3/1] | [2/3] |
| Number of circularized contigs | 3 | 1 | 0 |
| Contig N50 length (bp) | 4,904,000 | 3,527,000 | 3,160,000 |
| G+C content (%) | 52.42 | 43.25 | 46.61 |
| Number of genes | 5331 | 4908 | 4435 |
| Gene density (%) | 86.80 | 88.10 | 86.30 |
| Completeness (%) | 100.00 | 100.00 | 98.68 |
| Number of rRNAs | 21 | 28 | 27 |
| Number of tRNAs | 77 | 106 | 100 |

COG classification assigned 80.5% of the CDS into functional categories (Fig. 2). The distributions of COG functional categories in chromosomes of *P. denitrificans* CCAH7 and *R. marisflavi* CCAH4 were similar to that of the BGC-deficient *Escherichia coli* K-12 substrain MG1655. Amino acid transport and metabolism, Transcription, and Carbohydrate transport and metabolism were the most represented categories in these three genomes, but Cell cycle control, cell division, chromosome partitioning was among the least abundant in *P. denitrificans* and *E. coli*, and among the most abundant in *R. marisflavi*. The distribution of COG functional categories in *P. elyakovii* was notably different, with Signal transduction mechanisms, Amino acid transport and metabolism, and Cell wall/membrane/envelope biogenesis being the most represented. COG functional categories in plasmids of the three isolates were very uneven, potentially due to the highly variable molecule sizes (Supp. Fig. 1).

**FIG 2.**
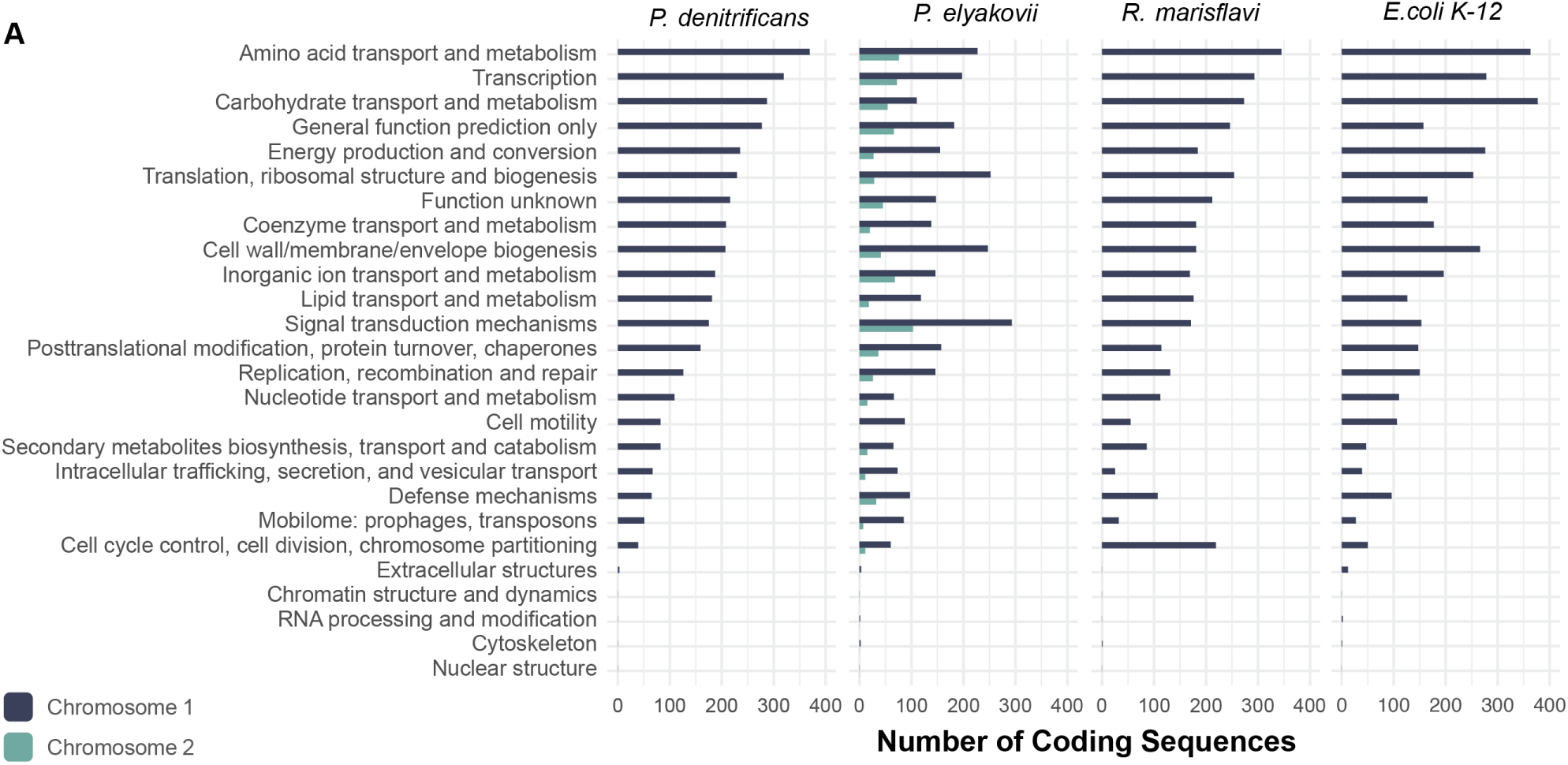
Representation of gene functions in genomes of the selected bacterial isolates and *Escherichia coli* strain K-12 substrain MG1655. Number of chromosomal protein-coding sequences (CDS) in functional categories determined by COGclassifier.

### Potential Genomic Basis of Antimicrobial Activity

KEGG metabolic pathway analysis revealed that the genome of *P. elyakovii* contains genes pertaining to *biosynthesis of siderophore group nonribosomal peptides*; within this pathway, all genes involved in the production of myxochelin A and B were present (Supp. Fig. 2). The genome of *R. marisflavi* contained *carotenoid biosynthesis* genes that suggest production of lycopene by this bacterium (Supp. Fig. 3). Also, the genomes of all three isolates encoded *streptomycin synthesis* genes, though no isolate contained all genes required for actual streptomycin synthesis (Supp. Fig. 4).

The genomes of the three isolates were further investigated to assess their biosynthetic potential in regard to secondary metabolite production. In total, 130 putative BGCs were predicted by several tools (see Methods) with different types of biosynthetic classifications. BGCs identified by antiSMASH were classified into 14 biosynthetic cores (Fig. 3A)(Tables S1, S2, S3), whereas BGCs identified by GECCO (Table S4) and deepBGC (Table S5) were classified into 5 biosynthetic classes, a few pertaining to one or more classes (Fig. 3B, C). Some of the more specific cores found by antiSMASH corresponded to 4 of the classes found by GECCO and deepBGC (Fig. 3). Five BGCs had considerable gene similarity (*≥*50%) to MiBiG BGCs implicated in the production of a range of metabolites, listed in Table 4.

**FIG 3.**
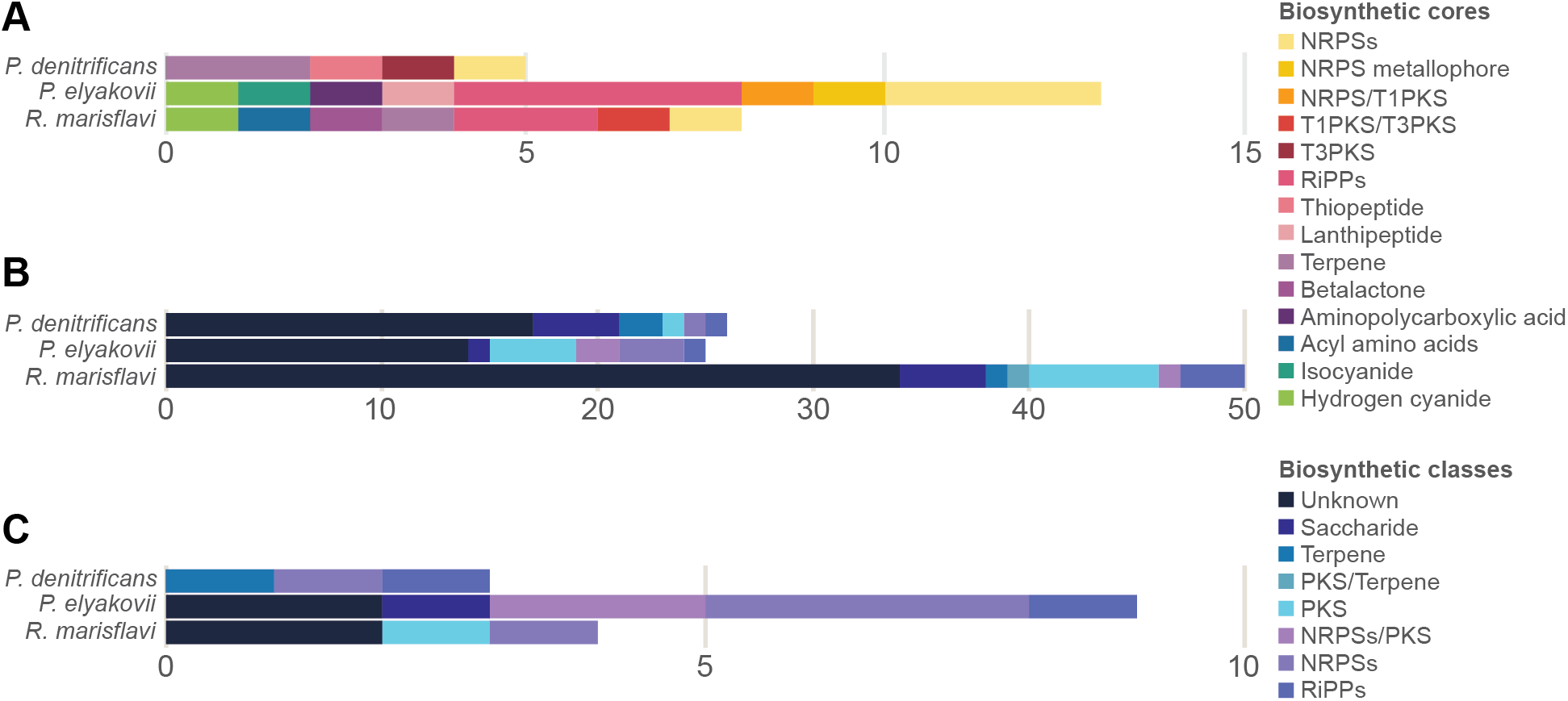
Counts of BGC cores and classes predicted for the three isolates using AntiSMASH **(A)**, DeepBGC **(B)**, and GECCO **(C)**.

**TABLE 4.** BGC hits with similarity to MiBiG BGCs and their respective products.

| **Isolate** | **MiBiG BGC Hits** | **Gene Similarity (%)** | **Product** |
| --- | --- | --- | --- |
| *P. denitrificans* | BGC0002123 | 100 | Pseudovibriamide A1-6, B1-6 |
| *P. elyakovii* | BGC0002491 | 100 | Myxochelin A, B, pseudochelin A |
| *P. elyakovii* | BGC0002345 | 66 | Hydrogen cyanide |
| *P. elyakovii* | BGC0000314 | 100 | Bromoalterochromide A |
| *R. marisflavi* | BGC0000645 | 50 | Carotenoid |

The *conservative* similarity network analysis revealed that most BGCs are unique with no similarity to reference clusters and that there was substantial redundancy between predictions from the different tools (Fig. 4A). Unique BGCs in the similarity network (i.e., singletons) encompassed 100 BGCs. Redundancy between predictions was showcased by twelve pairings that grouped BGCs predicted by different tools but with the same genomic location. Further, three clusters (each containing three BGCs), grouped a MiBiG reference BGC, and an antiSMASH and a GECCO prediction, the two latter also occurred at the same genomic location. While 14 BGCs grouped with MIBiG clusters in the *less-conservative network*, most BGCs were still classified as singletons, confirming the little representation of the identified BGCs in the reference database (Fig. 4B). Based on the network analyses and genomic locations of the predicted BGCs, we determined that a total of 115 non-redundant BGCs were predicted from these bacterial genomes: 55 from *P. denitrificans*, 33 from *P. elyakovii*, and 27 from *R. marisflavi*; no BGC was shared between isolate genomes.

**FIG 4.**
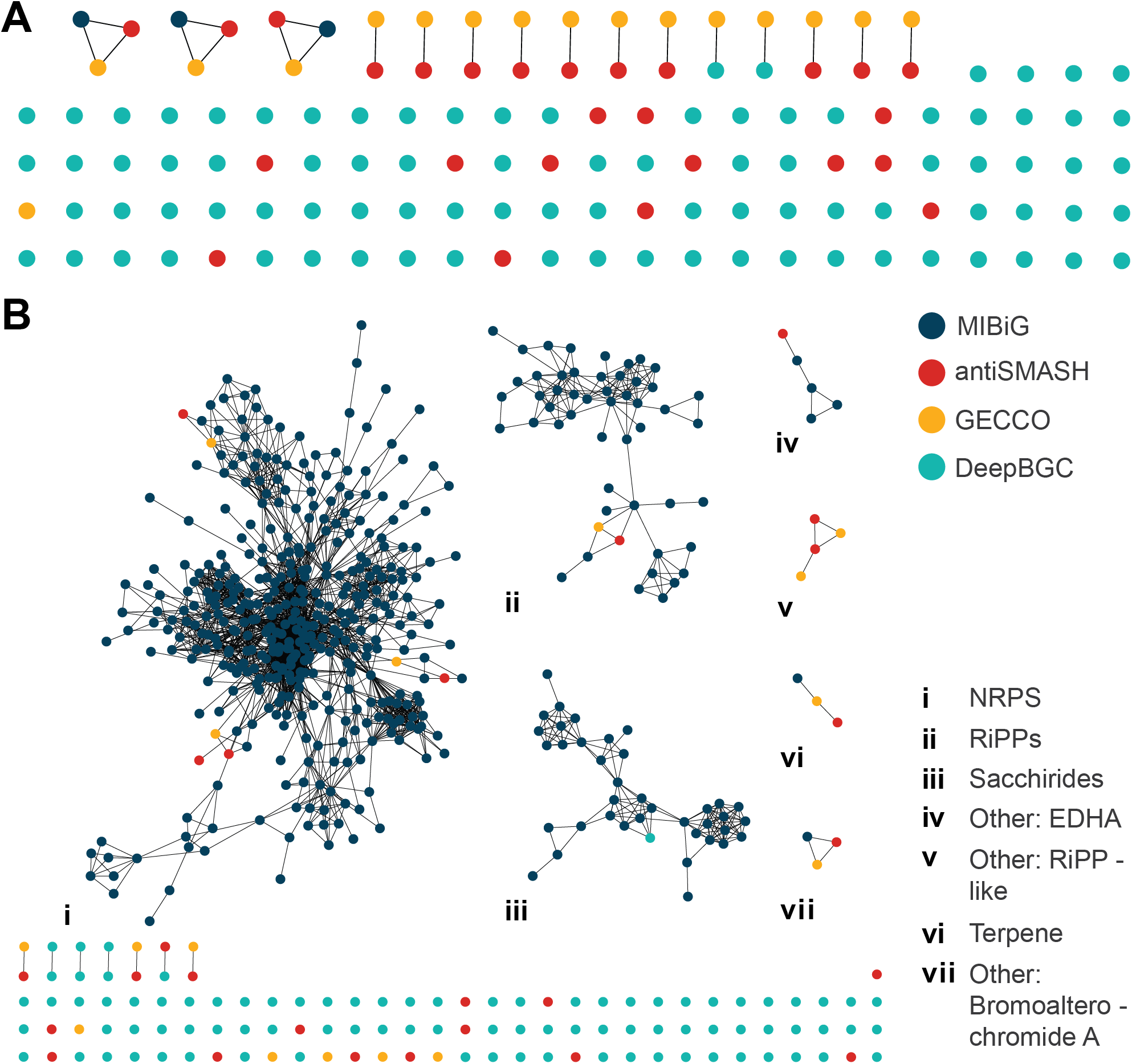
*Conservative* **(A)** and *less-conservative* **(B)** BGC similarity networks. Nodes represent individual BGCs, colored by the computational tool used for prediction (see bottom right legend). Edges indicate gene similarity (*≥* 70% for conservative and *≥*30% for less-conservative) between BGCs. Clusters are labeled (i-vii) according to their predicted product class following the bottom right legend.

## DISCUSSION

The present study investigated culturable microorganisms from the CCA *Hydrolithon boergesenii*. as a first bioprospecting step. Three of them exhibited antagonistic activity against several human and marine pathogens, some of which were resistant to common antimicrobials in the market. To further assess the antimicrobial potential of these three isolates, their genomes were sequenced. Genes and BGCs potentially underpinning their antimicrobial phenotype were identified. Below, we discuss how these genes and BGCs might be implicated in the antimicrobial activity of these bacteria and in other aspects of their ecology.

### Functional Roles of CCA Bacterial Isolates

The different gene functions and metabolic pathways found in each isolate (Fig. 2, Supp. Fig. 5) suggests that the isolates are occupying distinct niches in the CCA microenvironment. In previous work, *Pseudovibrio denitrificans* has been identified as an important stabilizing agent of reef ecosystems through its coral settlement induction properties [49][50], while simultaneously inhibiting larval settlement of other invertebrates, like barnacles and bryozoans [51]. Coral settlement and metamorphosis has been largely attributed to the secondary metabolite tetrabromopyrrole (TBP) [52].

Nevertheless, no genes (*bmp1-10* [41]) or BGCs involved in the generation of TBP could be found (Table 4). This observation indicates that: (i) this particular strain is not able of inducing coral larvae settlement, (ii) the biosynthetic pathways of TBP are not fully known yet, or (iii) other inducers yet to be fully characterized might be responsible for this trait [53][54]. Future research on this *P. denitrificans* strain should therefore focus on verifying whether this bacterium can produce TBP and whether it can induce coral settlement, which could lead to the description of new coral larvae settlement inducers.

Various members of the genus *Pseudoalteromonas* have also been shown to induce coral settlement using TBP as a chemical cue [21]. However, similar to *P. denitirifcans*, no genes involved in TBP production were found in the *Pseudoalteromonas elyakovii* strain isolated in this study. Beyond this, *P. elyakovii* has been previously found in the mussel *Crenomytilus grayanus* and in the brown seaweed *Laminaria japonica* [55]; this is the first instance, to our knowledge, that this bacterium has been isolated from a member of the phylum Rhodophyta. Within *Laminaria*, *P. elyakovii* has been isolated from spot-wounded fronds, suggesting that *P. elyakovii* may be causing damage to this alga. Further, our analyses of metabolic pathways (Supp. Fig. 2) and BGCs (Table 4) suggest that this isolate may be actively acquiring iron from the environment and/or host through siderophores like myxochelin A and B, enterochelin, and pseudochelin A. Iron acquisition mechanisms drive potential virulence of well-established pathogens [56][57]. This bacterium might thus be an algal pathogen.

The genome of the other bacterium recovered here, *Rossellomorea marisflavi*, contained genes implicated in the resistance to arsenic (i.e., *arsB*, *arsC*, and *arsR*) and in the production of the antioxidant carotenoid lycopene (Table 4 and Supp. Fig. 3). A genomic characterization of another member of the same genus, *Rossellomorea* sp. y25, found similar metabolic capabilities, with added resistance to cadmium and zinc [58]. These genetic features were hypothesized to play a role in the adaptation to deep-sea environments, where this bacterium was isolated from. In *R. marisflavi*, resistance to heavy metals might be associated with the potential for bioaccumulation of trace elements of CCA [59]. Though we are not aware of any reports of arsenic in these algae, contamination with heavy metals has occurred several times in Cartagena Bay [60][61][62][63], where the CCA analyzed here was sampled. Lycopene production, on the other hand, might be related to its ability to prevent the lethal action of UV radiation on bacterial cells [64] given the high-light environments where CCA inhabit.

### Antimicrobial Activity is Likely Due to BGCs with Unknown Functions

Metabolic pathway exploration showed no prominent representation of genes involved in antibiotic biosynthesis in either of the three CCA isolates (Supp. Figs. 6-8). Nevertheless, metabolic complementation [65] among members of the CCA microbial consortium might be a mechanism leading to the biosynthesis of some antimicrobials. For instance, while none of the three isolates possessed the complete pathway for biosynthesis of streptomycin, all of them contained several of the implicated genes (Supp. Fig. 4). However, antibiotic biosynthetic pathways are not likely to explain the antimicrobial activity observed in the three isolates.

Antimicrobial activity is often a result of products of BGCs. The most frequent BGC classes identified in this study included non-ribosomal peptides (NRPs), ribosomally synthesized and post-translationally modified peptides (RiPPs), polyketides (PKS), saccharides, and terpenes [23]. These biosynthetic classes are commonly identified in bioprospecting studies of prokaryotes in marine environments [66]. Nevertheless, the number of BGCs recovered in the genomes of the three isolates studied here was unusually high. For example, BGCs predicted by antiSMASH ranged between five (in *R. flavimaris*) and 13 (in *P. elyakovii*), whereas a former large-scale study predicted 18,043 BGCs from 5,743 microbial genomes (i.e., on average 3.14 BGCs per genome) [67].

Only five BGCs could be annotated with a known function (Table 4), from which three were predicted to biosynthesize known secondary metabolites. One of the BGCs, identified in the genome of *P. denitrificans*, is involved in the production of pseudovibriamide A1-6 and B1-6, which are hypothesized to play a role in surface motility [68]. The other two BGCs with known function were found in the genome of *P. elyakovii*. One of them is implicated in the production of catecholate-type siderophores, like myxochelin A and B, and pseudochelin A [69]. While these siderophores are crucial for bacterial iron homeostasis [69], they also exhibit weak activity against Gram-positive bacteria [70]; yet *P. elyakovii* inhibited both Gram-positive and -negative pathogens in antagonistic assays. The other BGC is associated with the production of bromoalterochromide A. *Pseuedoalteromonas* bromoalterochromides often have antibacterial activity [71], but no antimicrobial properties have been observed in bromoalterochromide A [72]. In addition, the carotenoid lycopene (Table 4 and Supp. Fig. 3) in *R. marisflavi* has been shown to have antimicrobial activity against Gram-negative bacteria, specifically against *Escherichia coli* [73] and members of the Chlamydiaceae family, such as *Chlamydia trachomatis* and *Chlamydia pneumoniae* [74]. While this carotenoid could explain some of the antimicrobial activity observed in the antagonistic assays, the *R. marisflavi* strain here did not inhibit the *E. coli* test strain, suggesting that *R. marisflavi* was not producing lycopene during the assays despite having the genomic capability. Therefore, the observed antimicrobial phenotypes are unlikely due to these BGCs.

Though biosynthetic pathways and known BGCs might underpin some of the antimicrobial activity revealed by the antagonistic assays, we hypothesize that this activity is most likely attributed to BGCs with unknown function. Given that most characterized biosynthetic pathways and BGCs have already been extensively explored for commercial antimicrobial production, these known compounds likely contribute to the existing resistance profiles of the tested pathogens. Hence, we believe that unknown metabolites are the ones to which the tested pathogens have not been commonly exposed to and are therefore susceptible to. BGCs can produce a wide range of metabolites [23], many of which remain to be explored; therefore, they could be responsible for this observed antimicrobial activity. Of special interest are the singletons, unique BGCs that appear only once in our dataset and show no significant similarity to known clusters. These singletons are particularly promising candidates for novel antimicrobial discovery as they represent previously uncharacterized biosynthetic targets.

### Future Directions

The advancement of bioprospecting efforts largely relies on deepening our understanding of fundamental microbial ecology. Certain microhabitats may be more prone to harboring novel antimicrobials, yet it remains difficult to identify which to target for bioprospecting. This knowledge gap extends beyond the ecology of microorganisms to the genomic basis of natural products, such as BGCs. Many BGCs remain uncharacterized and their prediction from genomic data is based on similarity to other known BGCs, which hinders bioprospecting efforts as this study supports.

One priority moving forward should be the consistent curation and functional characterization of unknown BGCs. To further the breadth of BGCs we identify, effort should be placed in employing multiple complementary approaches rather than relying on a single tool. This strategy allows for more comprehensive identification of BGCs that might be missed when using a single method. Furthermore, new methods that expedite the bioprospecting pipeline are also warranted. In this study, we explored several methods to identify BGCs of interest and demonstrated how network analyses can effectively prioritize promising BGCs by comparing their similarity to previously characterized clusters. This approach enables more targeted investigation of potentially novel compounds while reducing the resources spent on redundant or less promising BGCs. Finally, though costly, the development and enrichment of BGC databases and their continuous curation will provide an essential foundation for more targeted and efficient antimicrobial discovery efforts.

In summary, our study highlights the potential of crustose coralline algae (CCA) as a valuable target for antimicrobial discovery. We highlight the wide range of metabolic potential of CCA-associated bacteria through metabolic pathway analysis and the identification of 97 singleton BGCs. These singletons represent promising targets for unlocking novel biosynthetic potential in CCA-associated bacteria.

## Acknowledgments

This project was supported by an NSF IOS grant 2227068. DLL thanks his Linda Letawa Schobert Enrichment Scholarship and Eberly College of Science Anita M. Collins Research Scholarship. RAGP was supported by an Eberly Postdoctoral Research Fellowship awarded by the Eberly College of Science at Penn State University.

## Data availability

Raw sequencing data is available under the SRA accession xxxx and genome assembly and annotation under RefSeq accession xxxx.

## Author contribution

MM and KR initially conceptualized the project. DLL and RAGP designed the analyses to be performed. DLL and JSRT performed the sequencing run. KR, MWH, and EKJ conducted the antagonistic assays. DLL and RAGP wrote the manuscript. All authors reviewed, commented on, and contributed to the final version of the manuscript.

## Correspondence

The first author can be reached at. The last author can be reached at.

---

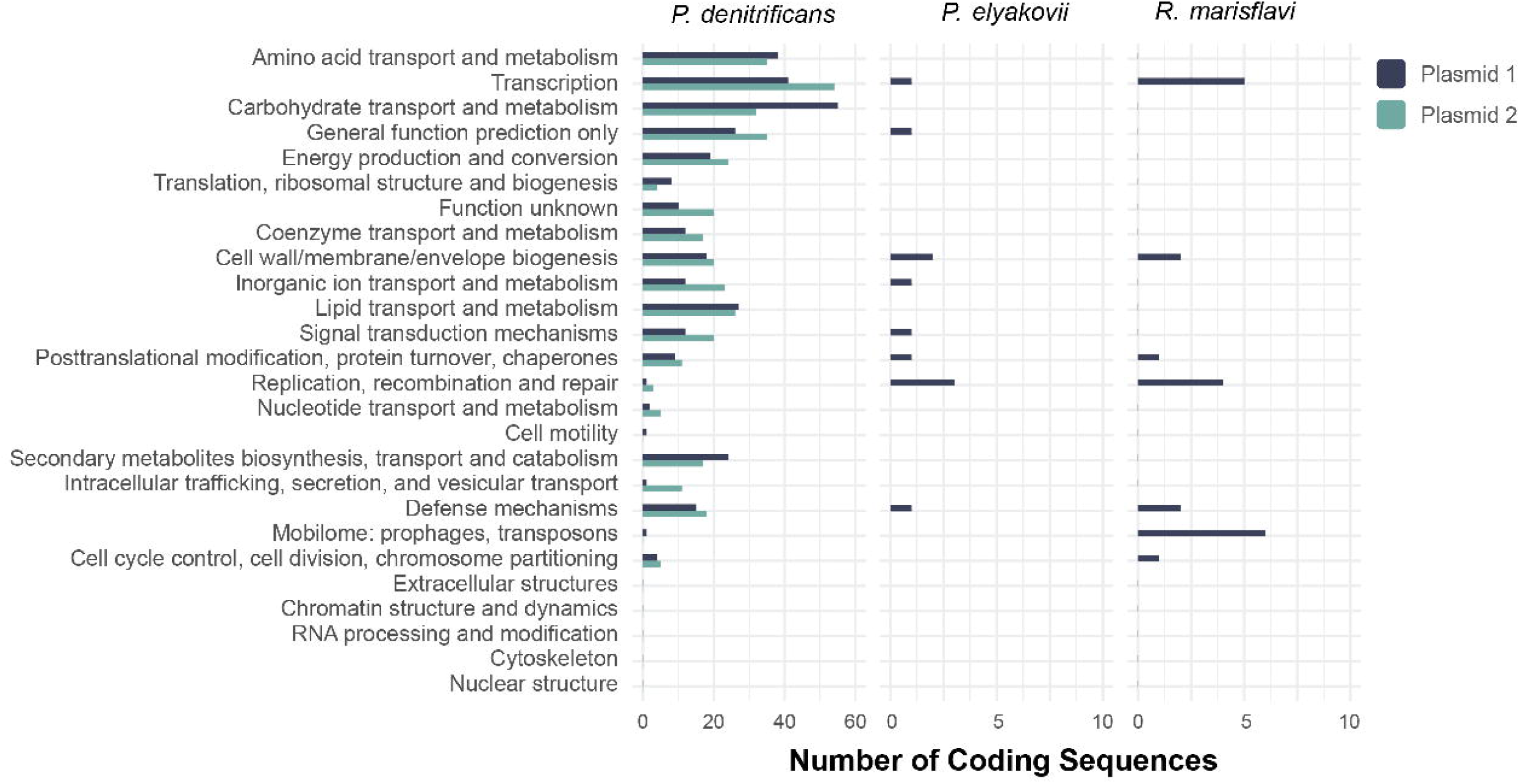

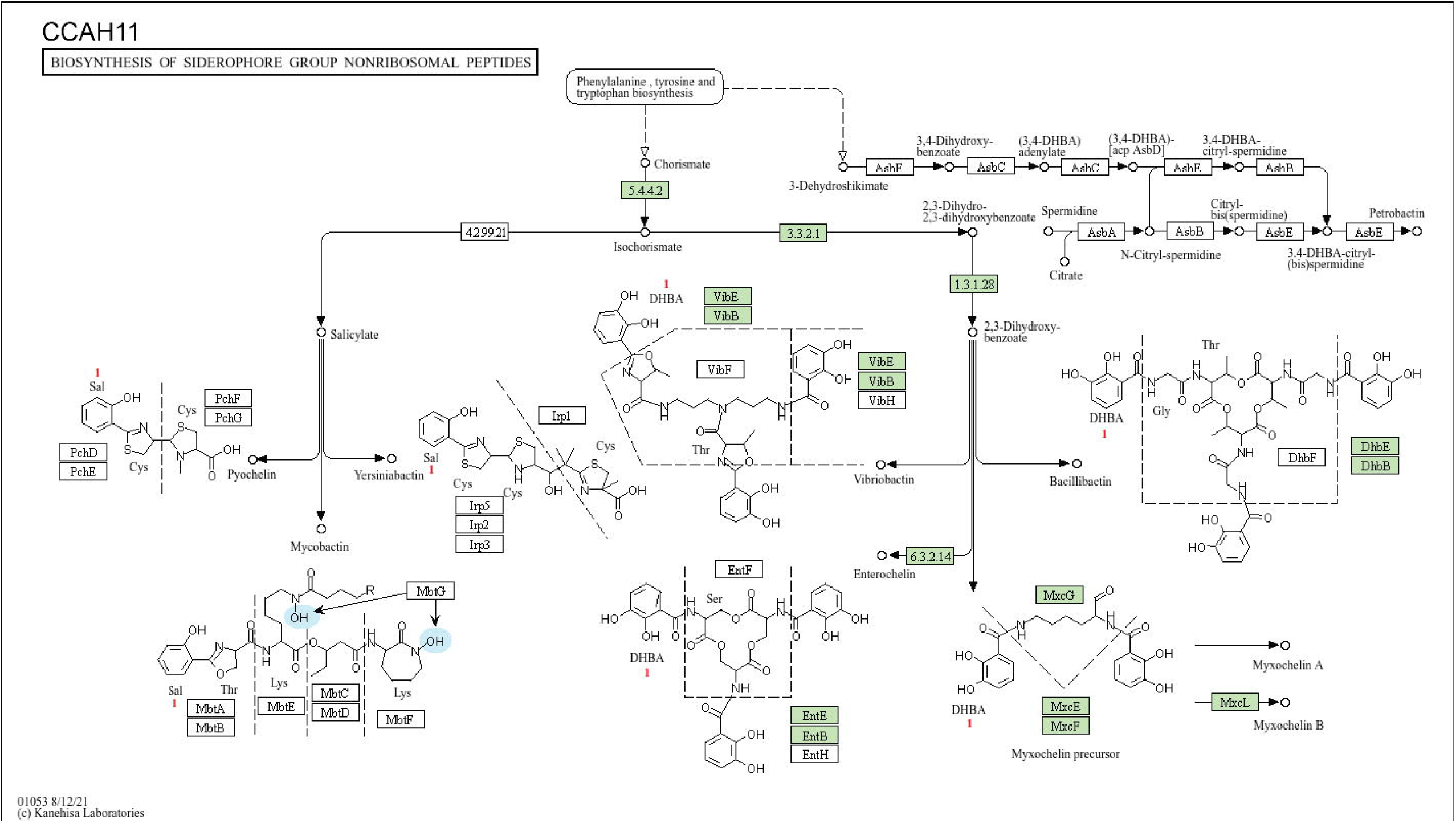

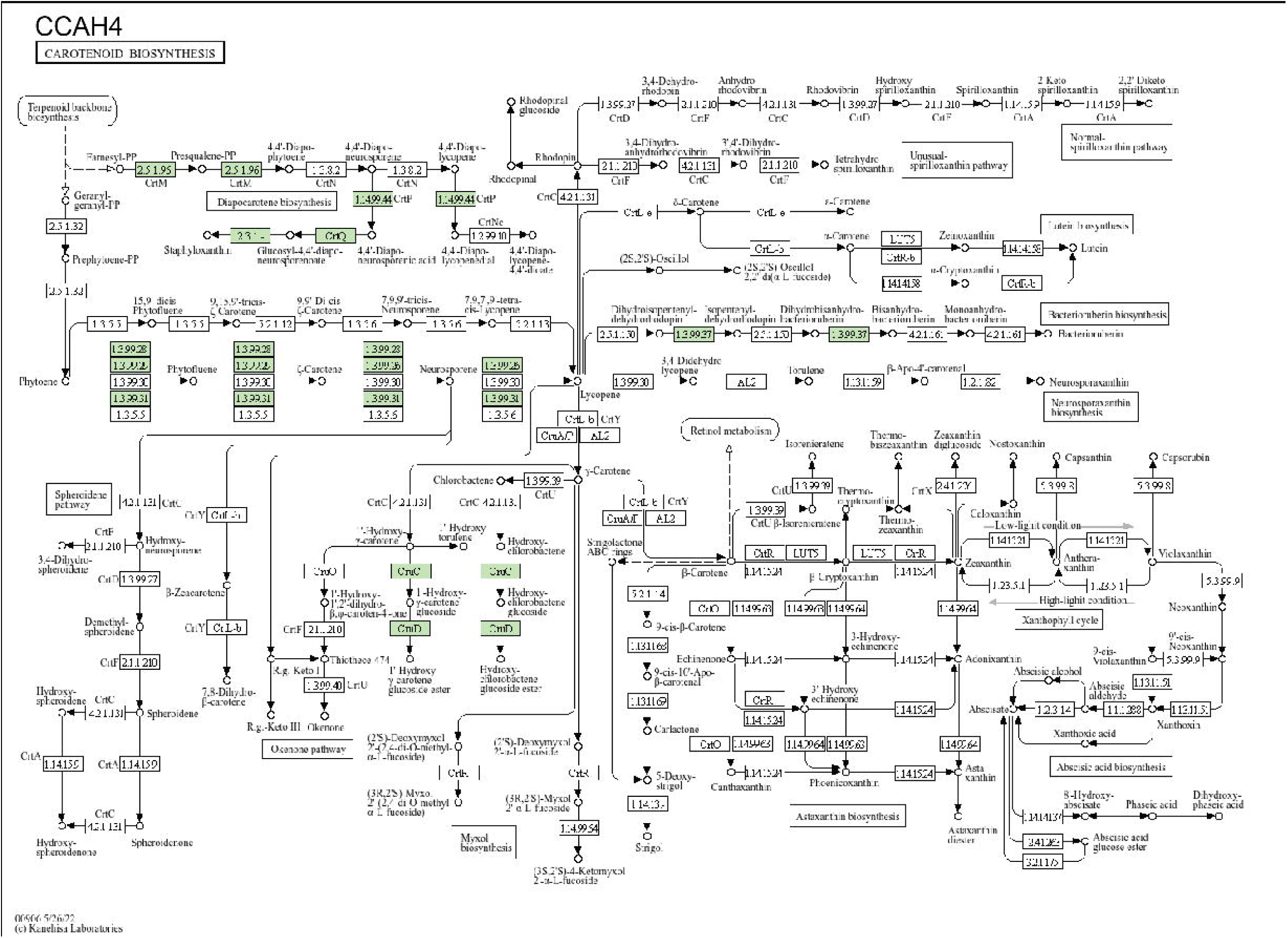

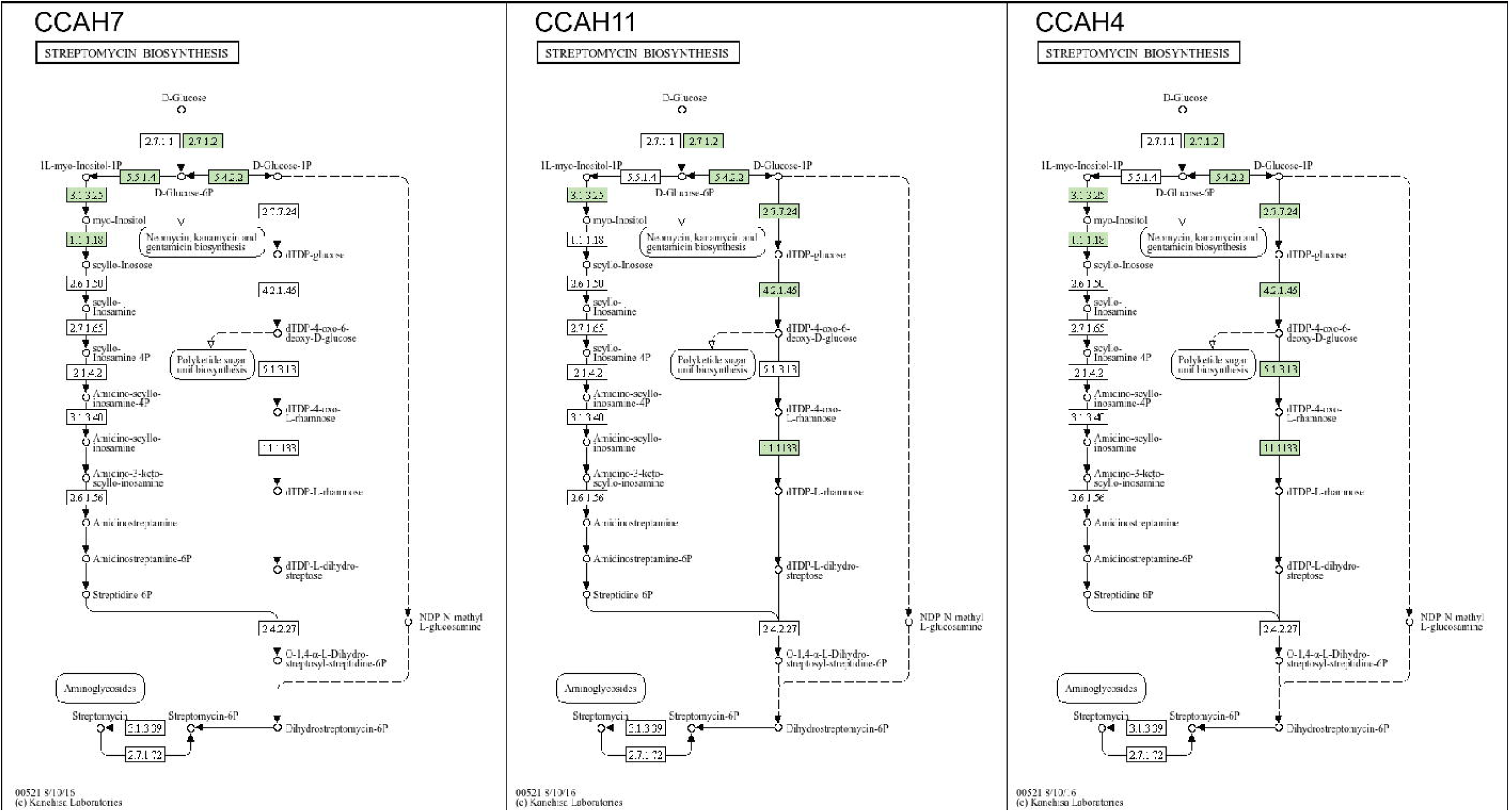

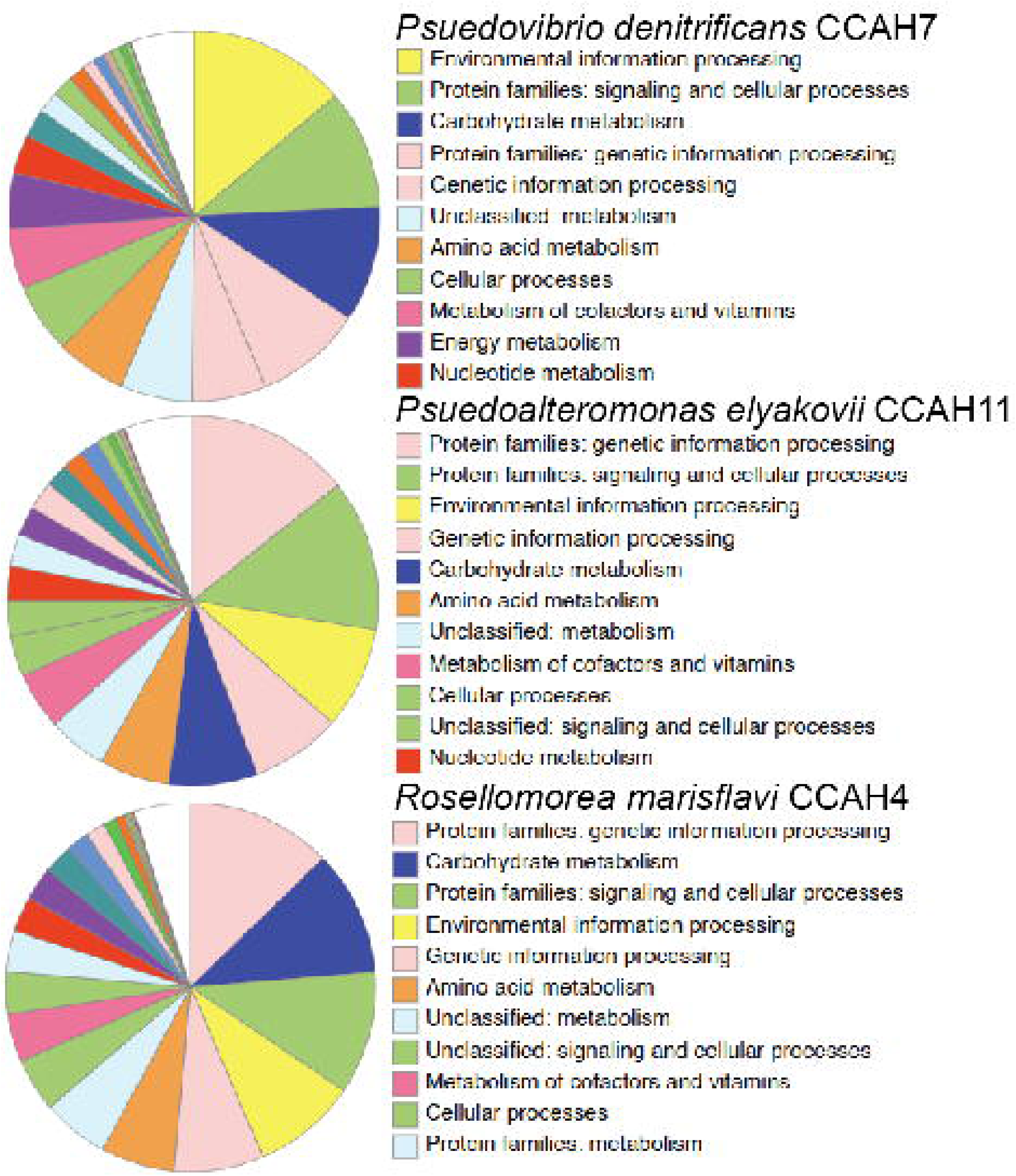

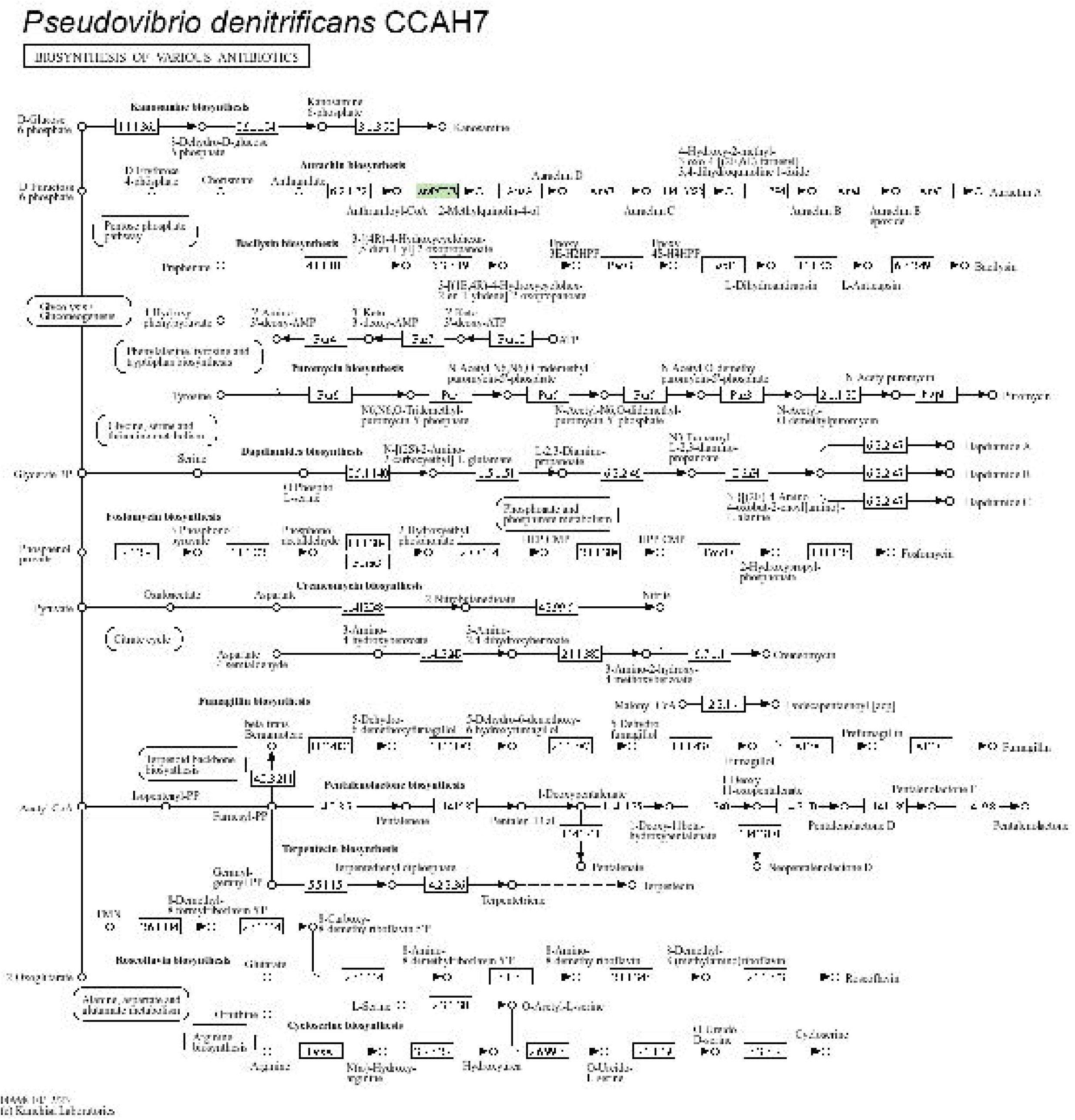

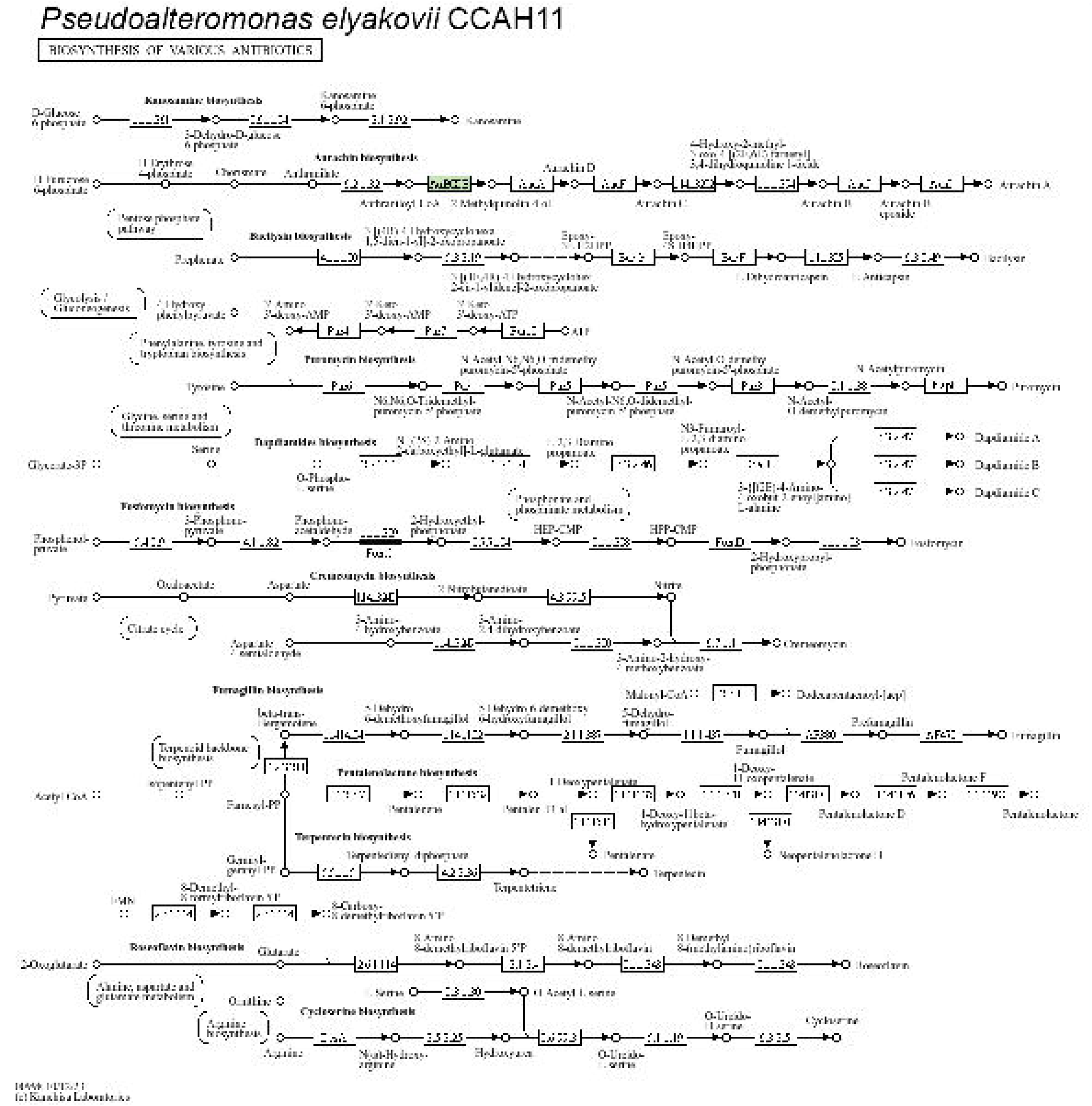

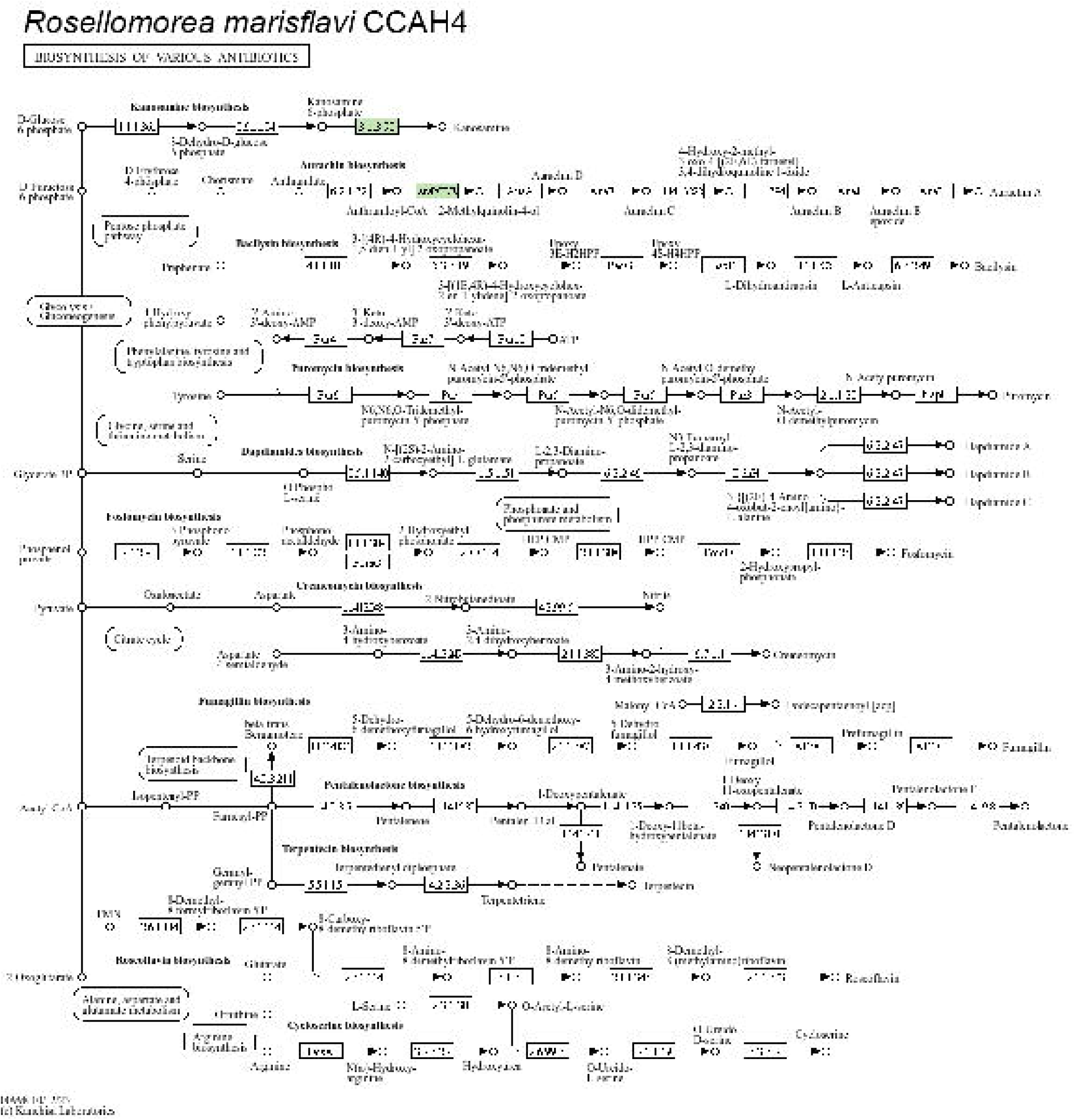

