## Supplementary Figures for "The Potential of CCA-associated Bacteria to Fight Antimicrobial-Resistant Pathogens: a Genomic Survey"

**TITLE:**

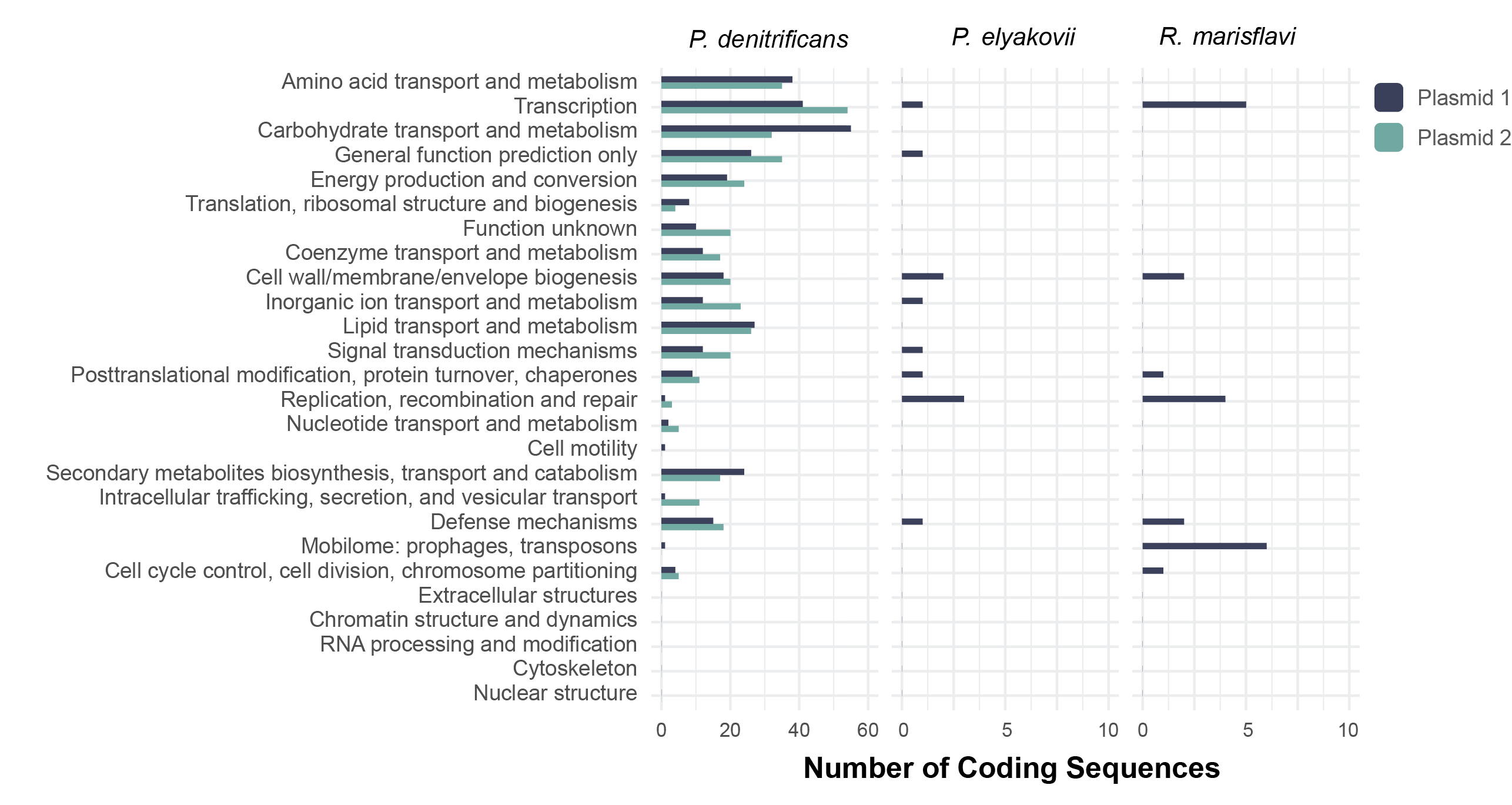

**Supplementary Figure 1.** Number of plasmid protein-coding sequences (CDS) in functional categories determined by COGclassifier. Note the different scales across panels.

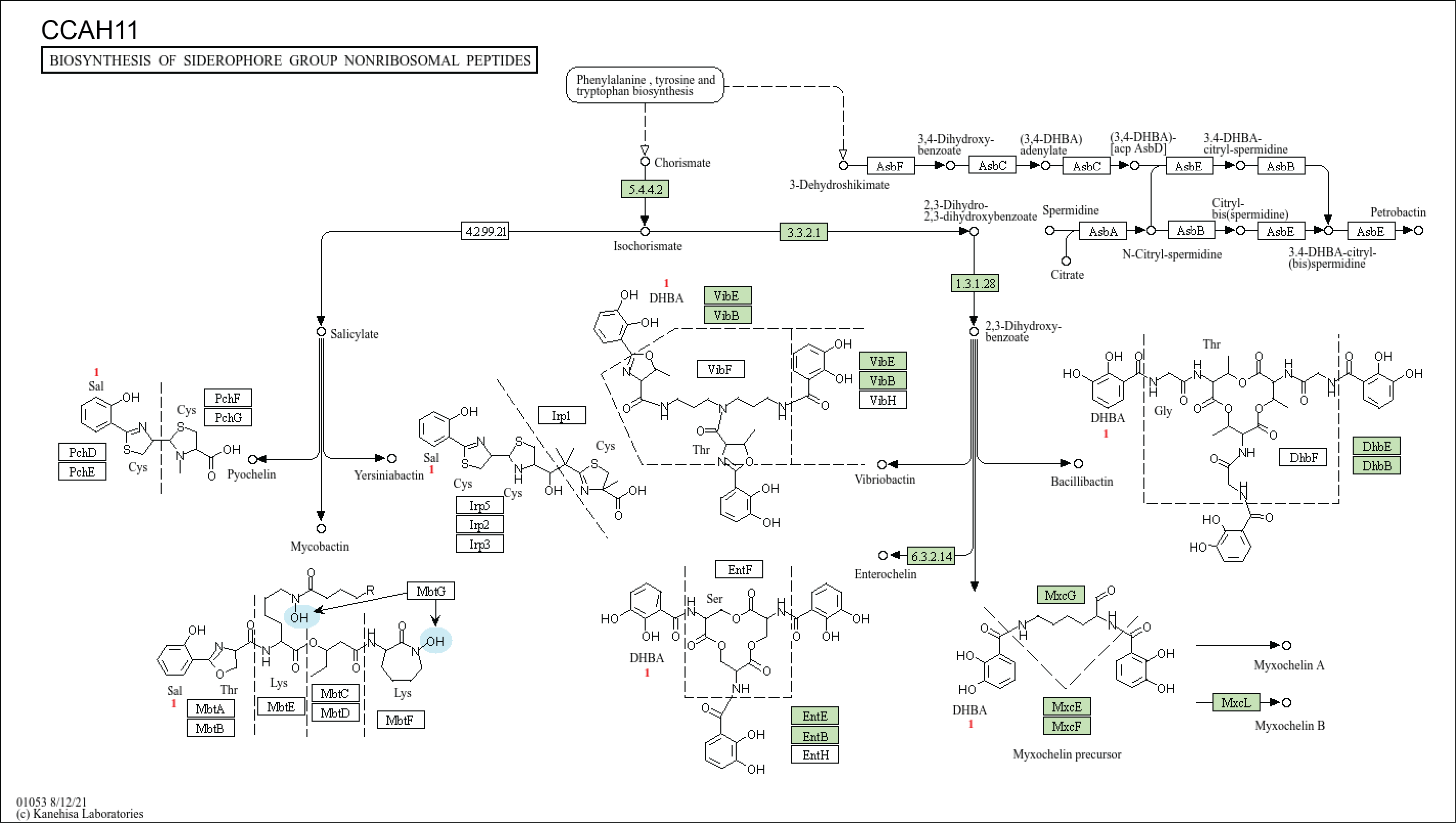

**Supplementary Figure 2.** KEGG pathway diagram of the biosynthesis of siderophore group nonribosomal peptides. Green boxes represent genes present in the genome of *Pseudoalteromonas elyakovii* CCAH11.

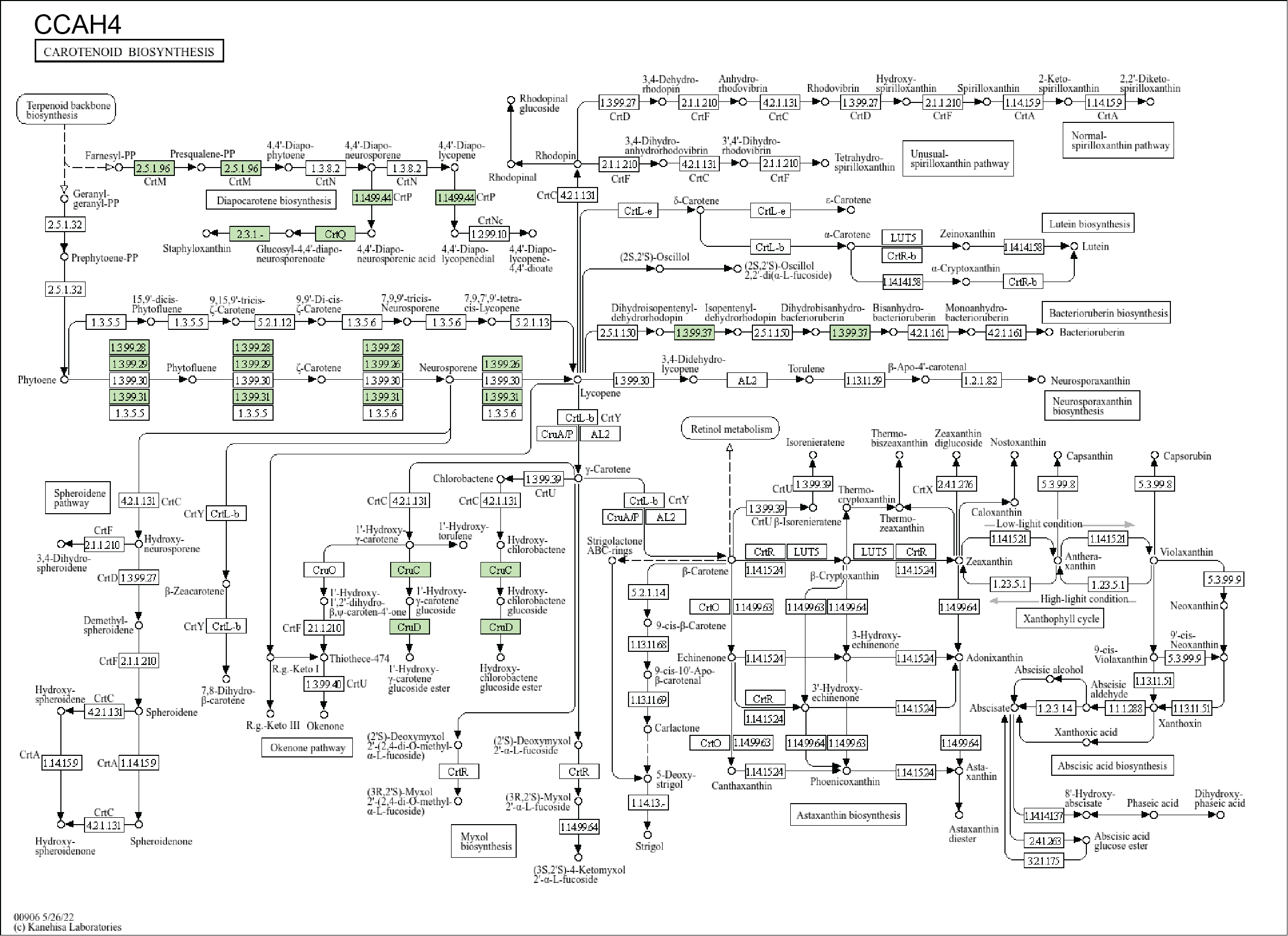

**Supplementary Figure 3.** KEGG pathway diagram of the biosynthesis of carotenoids. Green boxes represent genes present in the genome of *Rossellomorea marisflavi* CCAH4.

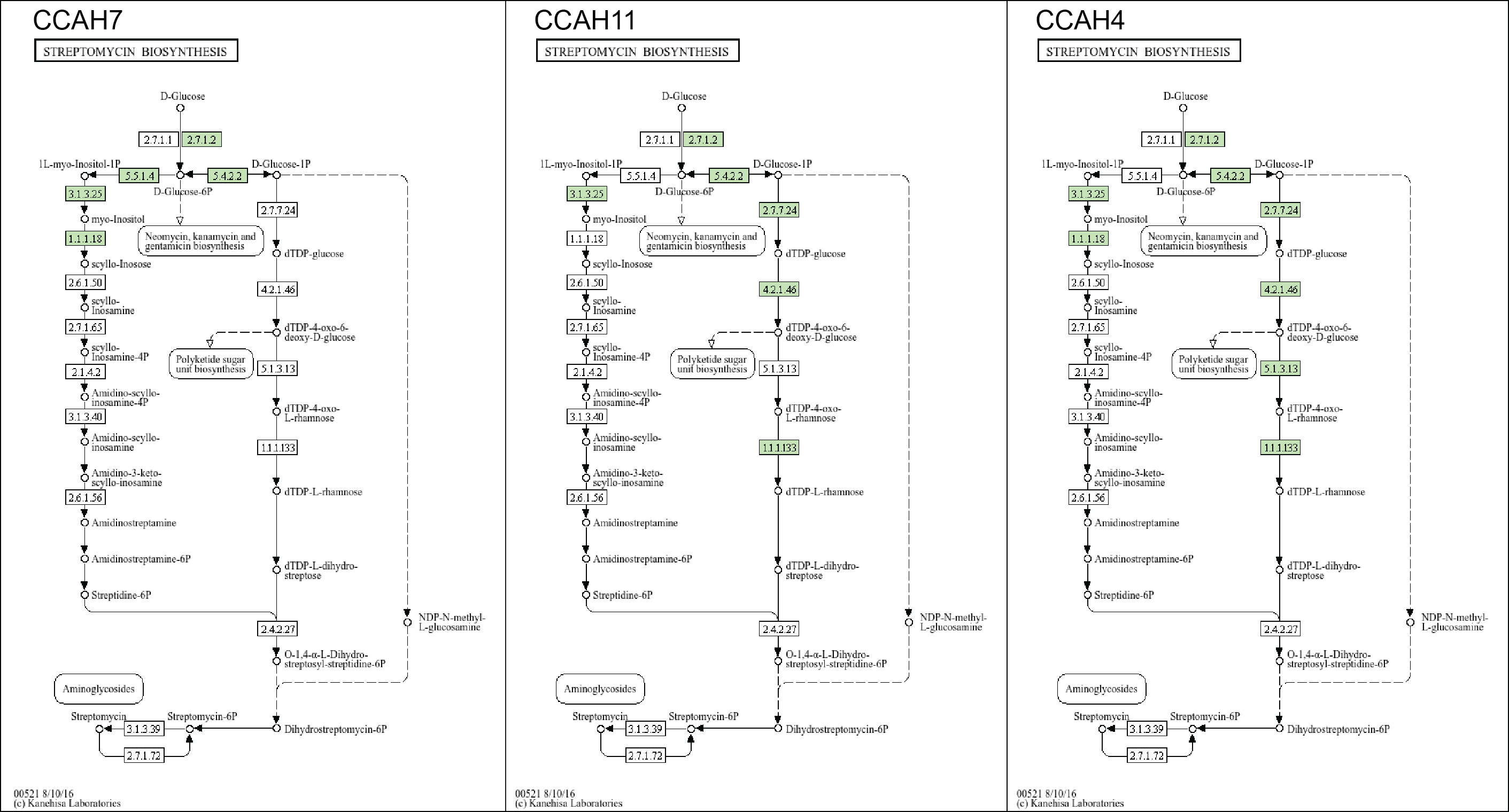

**Supplementary Figure 4.** KEGG pathway diagram of the biosynthesis of streptomycin. Green boxes represent genes present in the genomes of *Pseudovibrio denitrificans* CCAH7 (left), *Pseudoalteromonas elyakovii* CCAH11 (middle), and *Rossellomorea marisflavi* CCAH4.

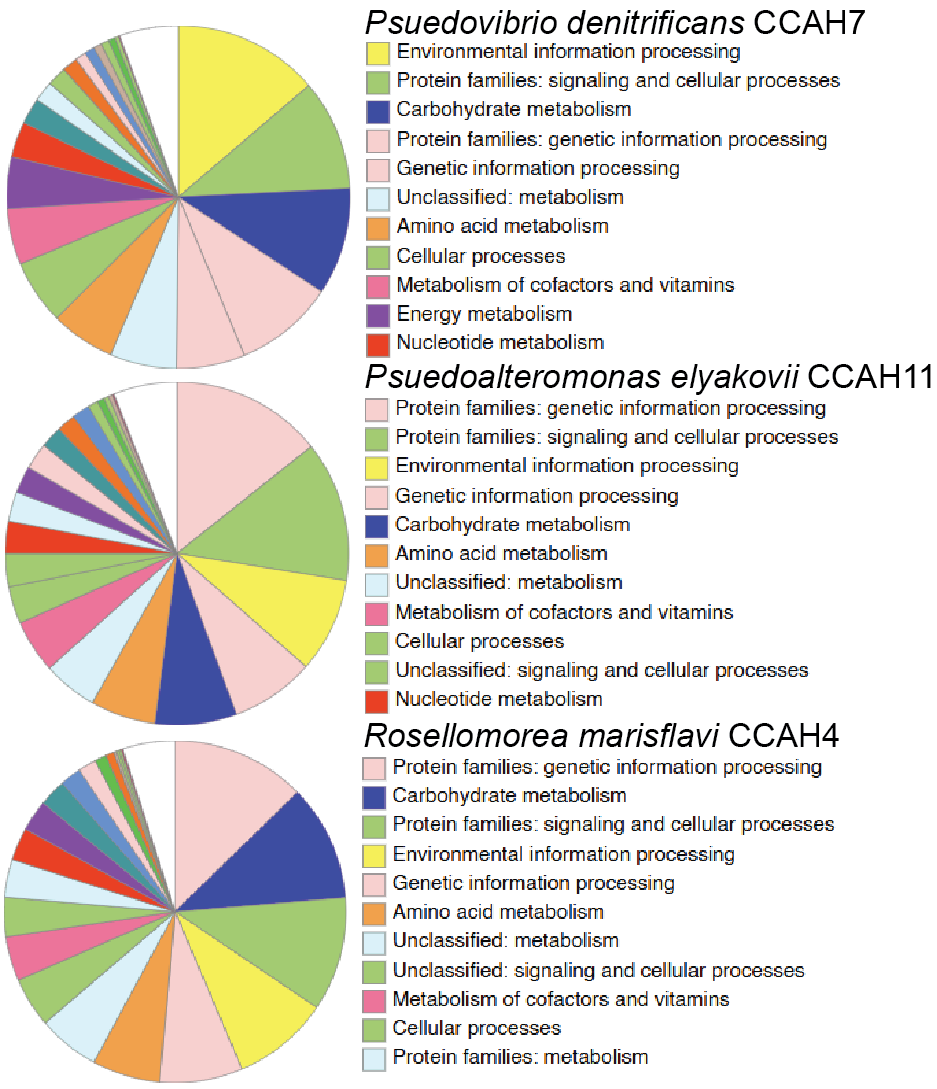

**Supplementary Figure 5.** Metabolic pathway representation in the genomes of *Pseudovibrio denitrificans* CCAH7 (top), *Pseudoalteromonas elyakovii* CCAH11 (middle), and *Rossellomorea marisflavi* CCAH4 (bottom), as determined by BlastKOALA annotation.

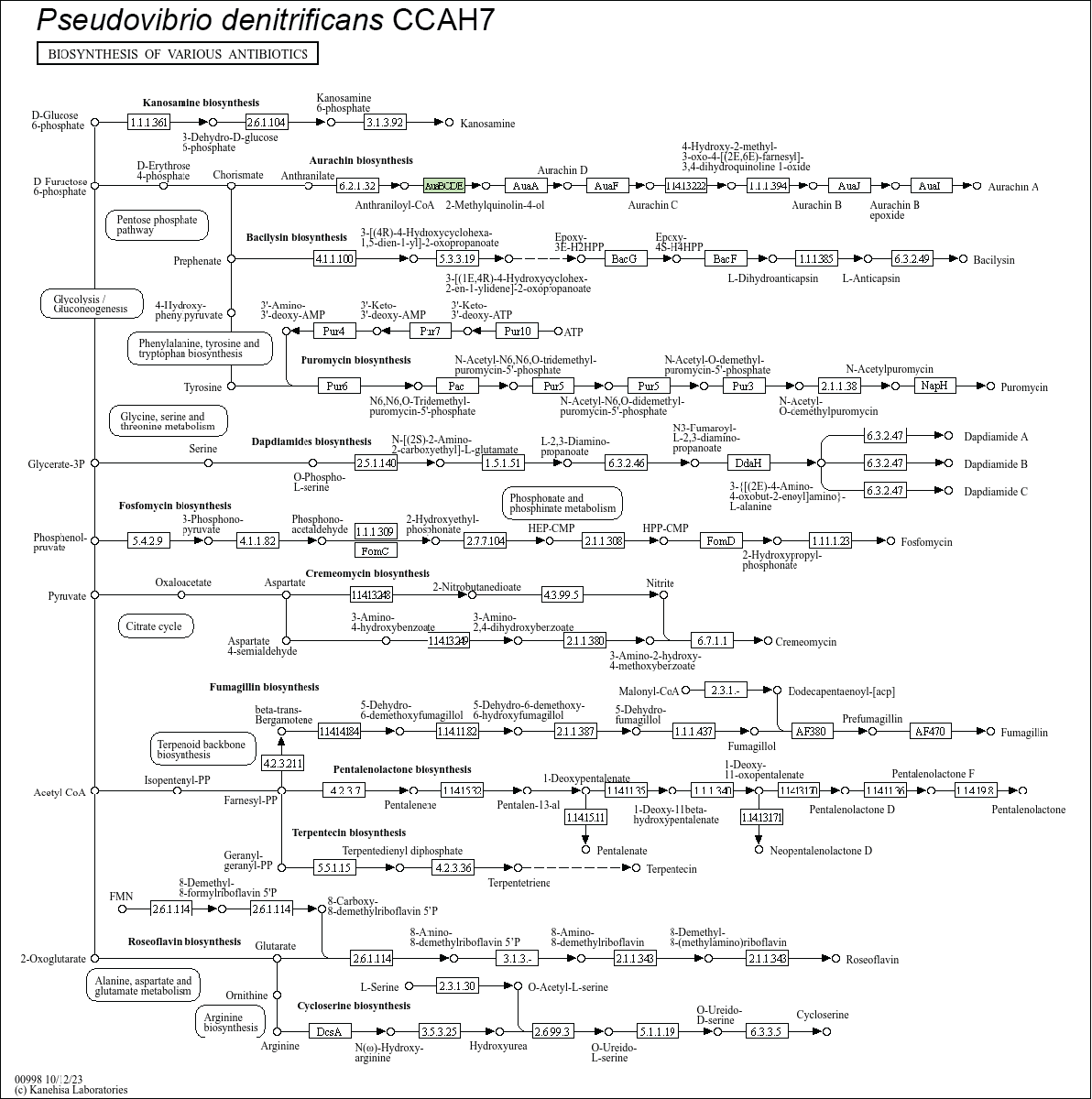

**Supplementary Figure 6.** KEGG pathway diagram of the biosynthesis of various antibiotics. Green boxes represent genes present in the genome of *Pseudovibrio denitrificans* CCAH7.

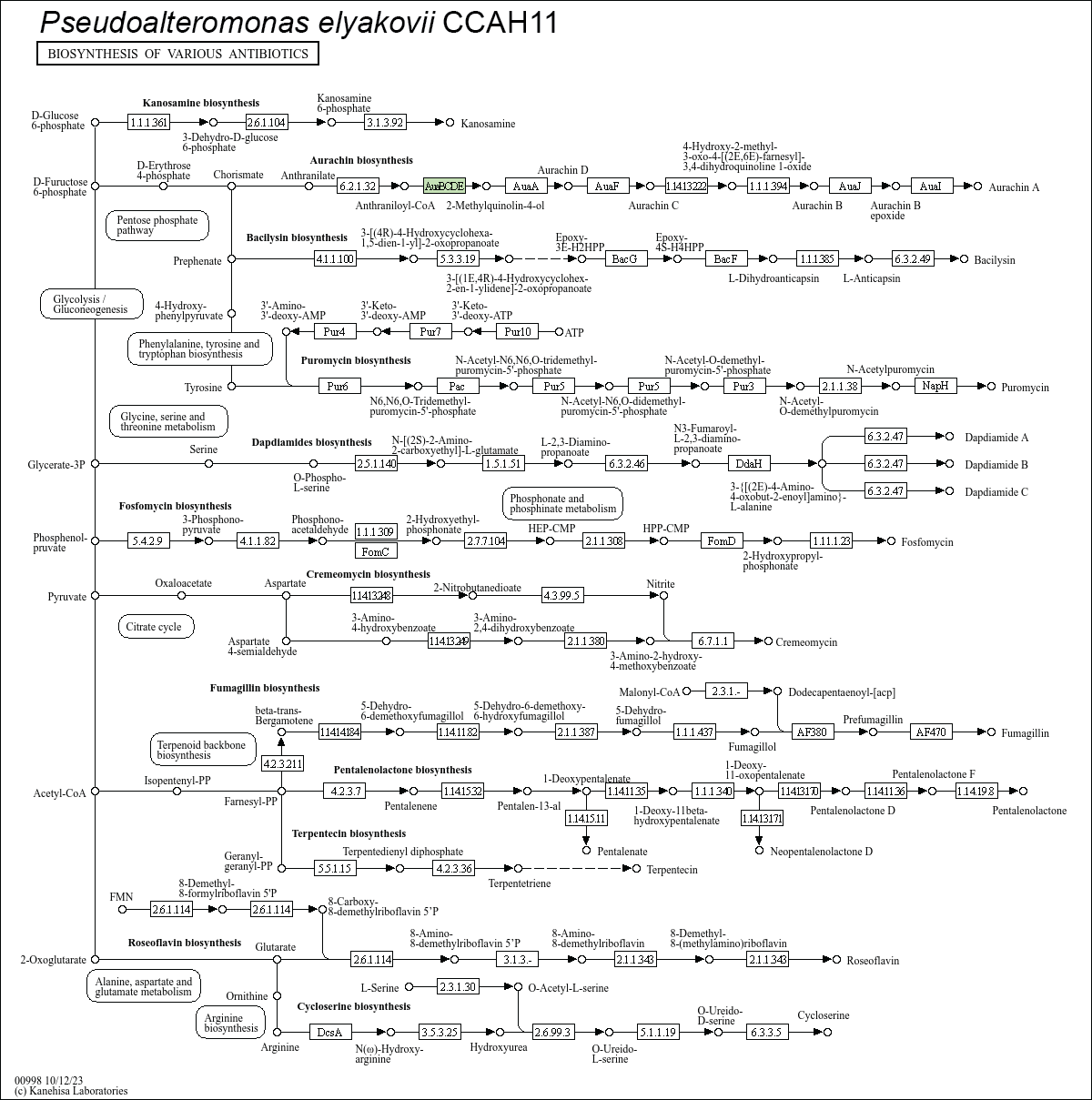

**Supplementary Figure 7.** KEGG pathway diagram of the biosynthesis of various antibiotics. Green boxes represent genes present in the genome of *Pseudoalteromonas elyakovii* CCAH11.

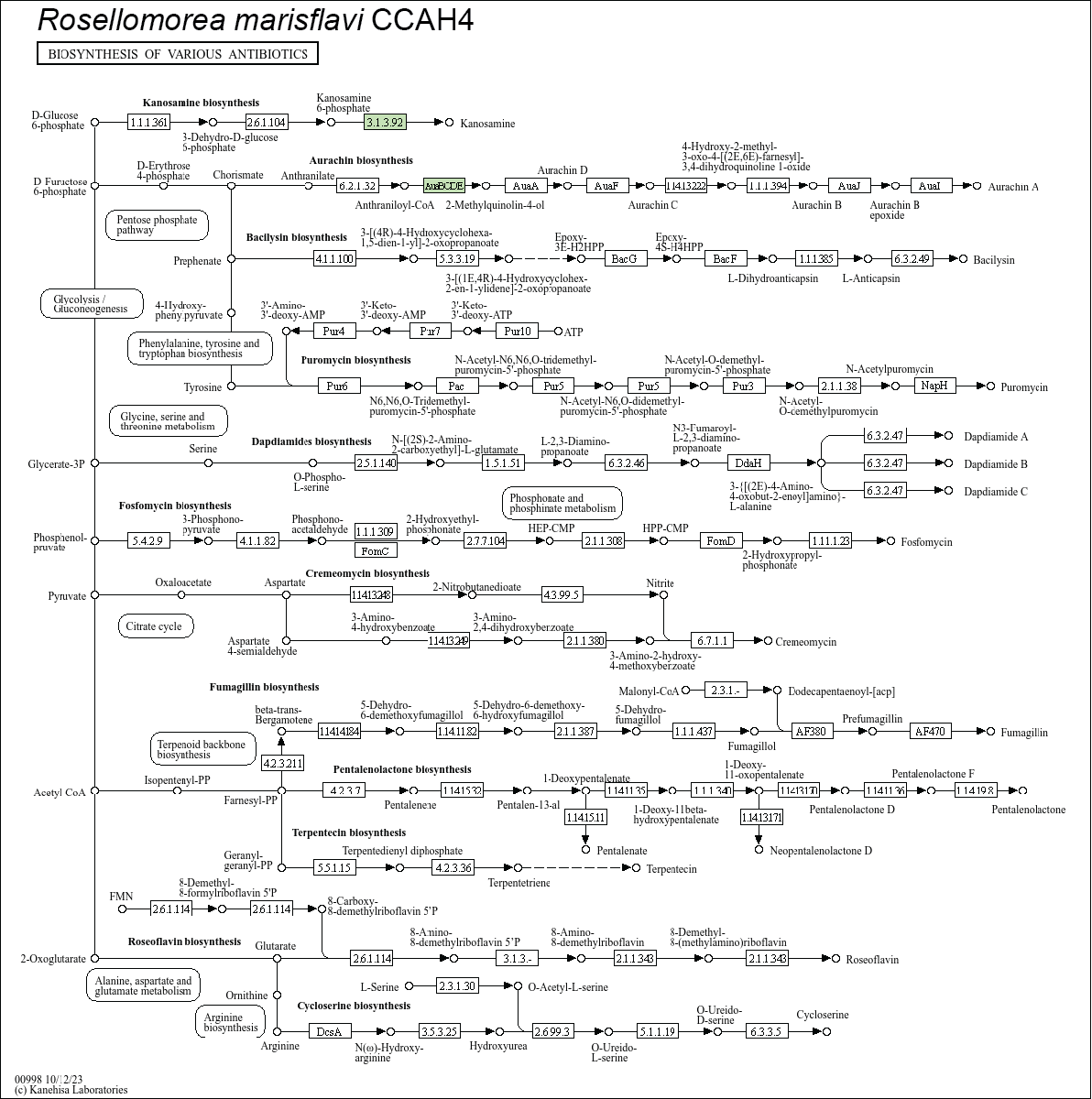

**Supplementary Figure 8.** KEGG pathway diagram of the biosynthesis of various antibiotics. Green boxes represent genes present in the genome of *Rossellomorea marisflavi* CCAH4.
